# Expansion of mutation-driven haematopoietic clones is associated with insulin resistance and low HDL-cholesterol in individuals with obesity

**DOI:** 10.1101/2021.05.12.443095

**Authors:** Rosanne C. van Deuren, Johanna C. Andersson-Assarsson, Felipe M. Kristensson, Marloes Steehouwer, Kajsa Sjöholm, Per-Arne Svensson, Marc Pieterse, Christian Gilissen, Magdalena Taube, Peter Jacobson, Rosie Perkins, Han G. Brunner, Mihai G. Netea, Markku Peltonen, Björn Carlsson, Alexander Hoischen, Lena M.S. Carlsson

**Author notes:** Corresponding authors* Corresponding authors: Alexander Hoischen, Departments of Human Genetics and Internal Medicine, Radboud Institute for Molecular Life Sciences, Radboud University Medical Center, Geert Grooteplein 10, 6525 GA Nijmegen, the Netherlands, Lena Carlsson, SOS Secretariat, Department of Molecular and Clinical Medicine, Vita Stråket 15, Sahlgrenska University Hospital, S-413 45 Gothenburg, Sweden. These authors contributed equally (Rosanne C. van Deuren, Johanna C. Andersson-Assarsson). These authors contributed equally (Alexander Hoischen, Lena M.S. Carlsson).

## Abstract

**Aims:** Haematopoietic clones caused by somatic mutations with ≥2% variant allele frequency (VAF), known as clonal haematopoiesis of indeterminate potential (CHIP), increase with age and have been linked to risk of haematological malignancies and cardiovascular disease. Recent observations suggest that smaller clones are also associated with adverse clinical outcomes. Our aims were to determine the prevalence of clonal haematopoiesis driven by clones of variable sizes, and to examine the development of clones over time in relation to age and metabolic dysregulation over up to 20 years in individuals with obesity.

**Methods and Results:** We used an ultrasensitive single-molecule molecular inversion probe sequencing assay to identify clonal haematopoiesis driver mutations (CHDMs) in blood samples from individuals with obesity from the Swedish Obese Subjects study. In a single-timepoint dataset with samples from 1050 individuals, we identified 273 candidate CHDMs in 216 individuals, with VAF ranging from 0.01% to 31.15% and CHDM prevalence and clone sizes increasing with age. Longitudinal analysis over 20 years in CHDM-positive samples from 40 individuals showed that small clones can grow over time and become CHIP. VAF increased on average by 7% (range -4% to 27%) per year. Rate of clone growth was positively associated with insulin resistance (R=0.40, P=0.025) and low circulating levels of high-density lipoprotein-cholesterol (HDL-C) (R=-0.68, P=1.74E^-05^).

**Conclusion:** Our results show that haematopoietic clones can be detected and monitored before they become CHIP and indicate that insulin resistance and low HDL-C, well-established cardiovascular risk factors, are associated with clonal expansion in individuals with obesity.

**Translational perspectives:** Clonal haematopoiesis-driver mutations are somatic mutations in haematopoietic stem cells that lead to clones detectable in peripheral blood. Haematopoietic clones with a variant allele frequency (VAF) ≥2%, known as clonal haematopoiesis of indeterminate potential (CHIP), are recognized as an independent cardiovascular risk factor. Here, we show that smaller clones are prevalent, and also correlate with age. Our longitudinal observations in individuals with obesity over 20 years showed that more than half of all clone-positive individuals show growing clones and clones with VAF <2% can grow and become CHIP. Importantly, clone growth was accelerated in individuals with insulin resistance and low high-density lipoprotein-cholesterol (HDL-C).

Translational outlook 1: Haematopoietic clones can be detected and monitored before they become CHIP.

Translational outlook 2: The association between insulin resistance and low HDL-C with growth of haematopoietic clones opens the possibility that treatments improving metabolism, such as weight loss, may reduce growth of clones and thereby cardiovascular risk.

**One Sentence Summary:** In obesity, the growth rate of mutation-driven haematopoietic clones increased with insulin resistance and low HDL-C, both known risk factors for cardiovascular disease.

## INTRODUCTION

Clonal haematopoiesis-driver mutations (CHDM) are somatic mutations that occur in haematopoietic stem cells (HSC) and lead to clones detectable in peripheral blood cells.^1, 2^ CHDMs have been described in several genes, with *DNMT3A, TET2, ASXL1*, and *JAK2* amongst the most frequently mutated.^1, 2^ The accumulation of somatic mutations is a hallmark of ageing^1^ and the prevalence of CHDMs increases with age^3^. Some studies suggest that clonal haematopoiesis may also be influenced by environmental stressors such as inflammation^1^, and that cellular stressors may promote the expansion of mutant clones^4^.

Individuals with CHDMs have increased risk of haematological cancers, but many CHDM carriers present without haematological diseases^1, 2^. Clonal haematopoiesis of indeterminate potential (CHIP) is defined as the state when the variant allele fraction (VAF) of CHDMs in peripheral blood DNA is ≥2%^5^. The presence of CHIP has been associated with increased all-cause mortality and risk of several non-haematological diseases, including myocardial infarction, heart failure and type 2 diabetes^2, 3, 6, 7^, and mechanistic studies in mice suggest that the relationship is causal^7-9^.

The risk of cardiometabolic diseases increases with age^10^, an association that may be strengthened by obesity. Obesity accelerates the normal ageing process, reducing life expectancy by 5 to 20 years^11-14^. Recent work suggests that CHDM prevalence in otherwise healthy older women is higher in those with obesity compared to those with normal body weight^15^, supporting the idea that obesity accelerates the age-related increase in clonal haematopoiesis. The mechanisms that link increased prevalence of CHDMs and obesity are unknown, but metabolic abnormalities (*e*.*g*. dyslipidaemia and glucose dysregulation) that are associated with obesity, cardiovascular disease and CHIP could play a role.

Clone size appears to correlate to risk of disease^7, 16^, with higher risk of both cardiovascular events and haematological malignancies in those with clones with VAF >10% compared to those with smaller clones^3^. The 2% VAF cut-off for CHIP clones is arbitrary and was chosen largely because of earlier technical limitations. However, a recent study identified a lower VAF threshold that is associated with worse outcome in patients with heart failure^17^. Unresolved questions include additional health consequences of smaller clones and how they evolve over time. In addition, the drivers of clonal outgrowth over time are largely unknown, and novel, potentially environmental, triggers may be involved. Furthermore, although most clones appear to be stable over time in healthy individuals^18^, growth of individual clones has not been investigated in individuals with obesity.

Here we used an ultrasensitive assay to analyse clonal haematopoiesis in blood samples from individuals with obesity from the longitudinal Swedish Obese Subjects (SOS) study. Our aims were to determine the prevalence of clonal haematopoiesis driven by clones of variable sizes, and to examine the development of clones over time in relation to age and metabolic dysregulation over up to 20 years in individuals with obesity.

## MATERIALS AND METHODS

### Subjects and samples

The present study includes participants from the control group of the SOS study.

Inclusion criteria were age between 37 and 60 years and body-mass index (BMI) of ≥34 for men and ≥38 for women; participants were recruited between September 1, 1987, and January 31, 2001, as previously described^12^. Details are given in Supplementary material online, *File S1*. Seven regional ethics review boards approved the study protocol and written or oral informed content was obtained. The study is registered at ClinicalTrials.gov (NCT01479452). All clinical investigations were conducted according to the principles expressed in the Declaration of Helsinki.

Two datasets were analysed from the individuals given usual care: a large single-timepoint dataset and a smaller multiple-timepoint dataset with up to 5 timepoints over 20 years. For the single-timepoint dataset, DNA was extracted from blood samples taken at a single timepoint for 1746 individuals. Of these, we excluded 668 samples because they did not pass DNA quality control. Thus, 1078 DNA samples of sufficient quality and quantity for sequencing were available, and from these we obtained high-quality sequencing data for 1050 samples. To analyse CHDMs over time, we selected 40 individuals for whom at least one CHDM had been detected in the single-timepoint dataset and blood samples for DNA extraction were available from baseline and up to four additional timepoints. The resulting multiple-timepoint dataset consisted of 180 DNA samples taken at baseline and at the 2-, 10-, 15-, and/or 20-year examinations (n=40, 38, 40, 38, and 24, respectively), and we obtained high-quality sequencing data for all 180 individual timepoint samples.

### CHDM detection by single-molecule molecular inversion probe (smMIP) sequencing

CHDMs were analysed by ultra-sensitive and error-correction based targeted sequencing, as previously described with minor modifications^19^ (Supplementary material online, *Files S1 and S2*). Briefly, smMIP sequencing of the entire *DNMT3A* gene as well as previously established CH-related hotspots in 23 additional genes (Supplementary material online, *Files S1 and S3)* was performed with two PCR and sequencing replicates for each DNA sample followed by two independent data-processing strategies and a targeted quality control to ensure high-quality. The average coverage over the entire CH-smMIP panel for each of two technical replicates per sample was 2840x and 3891x for our single- and multiple-timepoint datasets, respectively (Supplementary material online, *File S4*).

### CHDM definitions and classifications

CHDMs were subclassified, based on the arbitrary VAF threshold to define CHIP, into small (VAF <2%) and large (≥2%) clones^5^.

In our multiple-timepoint dataset, we classified CHDMs based on their evolvement during follow-up. CHDMs present in at least three timepoints for an individual were classified as traceable trajectories, whereas CHDMs only observed at a single or two timepoints were classified as events. Traceable trajectories were then further subclassified based on clone dynamics: 1) growing trajectories – where the absolute VAF of CHDMs at the final timepoint was at least 0.5% higher than at the first timepoint, 2) shrinking trajectories – where the VAF of CHDMs at the final timepoint was at least 0.5% lower than at the first timepoint, and 3) static trajectories – where the VAF of CHDMs at the final and first timepoints differed by less than 0.5%. As our trajectory definition requires CHDMs to be present at three or more timepoints, we annotated events with a VAF ≥2% at the final timepoint as ‘late-appearing clones’, as the abrupt appearance of a CHDM with such a high VAF likely suggests a fast-growing clone and therefore has the potential to be a growing trajectory. Finally, we computed relative VAF metrics that define CHDM evolvement over time by comparing the VAF at each timepoint to the first timepoint at which the clone was detected.

### Statistical analyses

In our cross-sectional, single-timepoint dataset, age differences were assessed using Wilcoxon-rank sum tests. We used logistic regression models to determine the impact of age on CHDM prevalence, and a linear regression model for the effect of age on CHDM size (log-transformed VAF).

Our longitudinal, multiple-timepoint dataset enabled us to determine the effect of age on clone growth, as dependent measurements allowed for tracing a single clone over time. To this end, we first selected a single CHDM trajectory per individual. We hypothesized that trajectories with higher VAFs are more important than trajectories with lower VAFs, since literature indicates that clone size correlates to the risk of disease^7, 16^. As such, we prioritized growing or shrinking trajectories over static trajectories, and for each individual selected the trajectory with the highest VAF detected at any follow-up timepoint as the ‘most important’ trajectory. Individuals in which the most important trajectory was shrinking were excluded from the subsequent analyses. We used a mixed linear model with random intercept and random slope, allowing varying effects of age per individual trajectory, to determine the effect of age on growth (VAF at each timepoint). To assess potential factors that could underlie differences in effect size of age on rate of growth, we evaluated the role of parameters related to glucose metabolism (glucose, insulin, homeostatic model assessment for insulin resistance (HOMA)-index), lipid metabolism (high-density lipoprotein cholesterol (HDL-C)), HSC proliferation, high-sensitivity C-reactive protein (hsCRP), and obesity in general (systolic blood pressure (SBP), body mass index (BMI), and waist-hip-ratio). We finally correlated the resulting random Beta estimates for age to clinical parameters averaged over the first three follow-up timepoints, by means of Spearman correlation coefficients, and subsequently expanded the mixed linear model with significant averaged clinical parameters. All statistical analyses were performed in R version 3.6.1 (R Core Team, URL https://www.R-project.org/), p-values <0.05 were considered statistically significant unless otherwise specified. For more details on specific analyses see Supplementary material online, *File S1*.

## RESULTS

### Subjects with available sequencing data

Using the ultra-sensitive smMIP assay (Figure 1A) we obtained high-quality sequencing data for 1050 individuals of our single-timepoint dataset and all 180 individual timepoints from our 20-year multiple-timepoint dataset.

**Figure 1.**
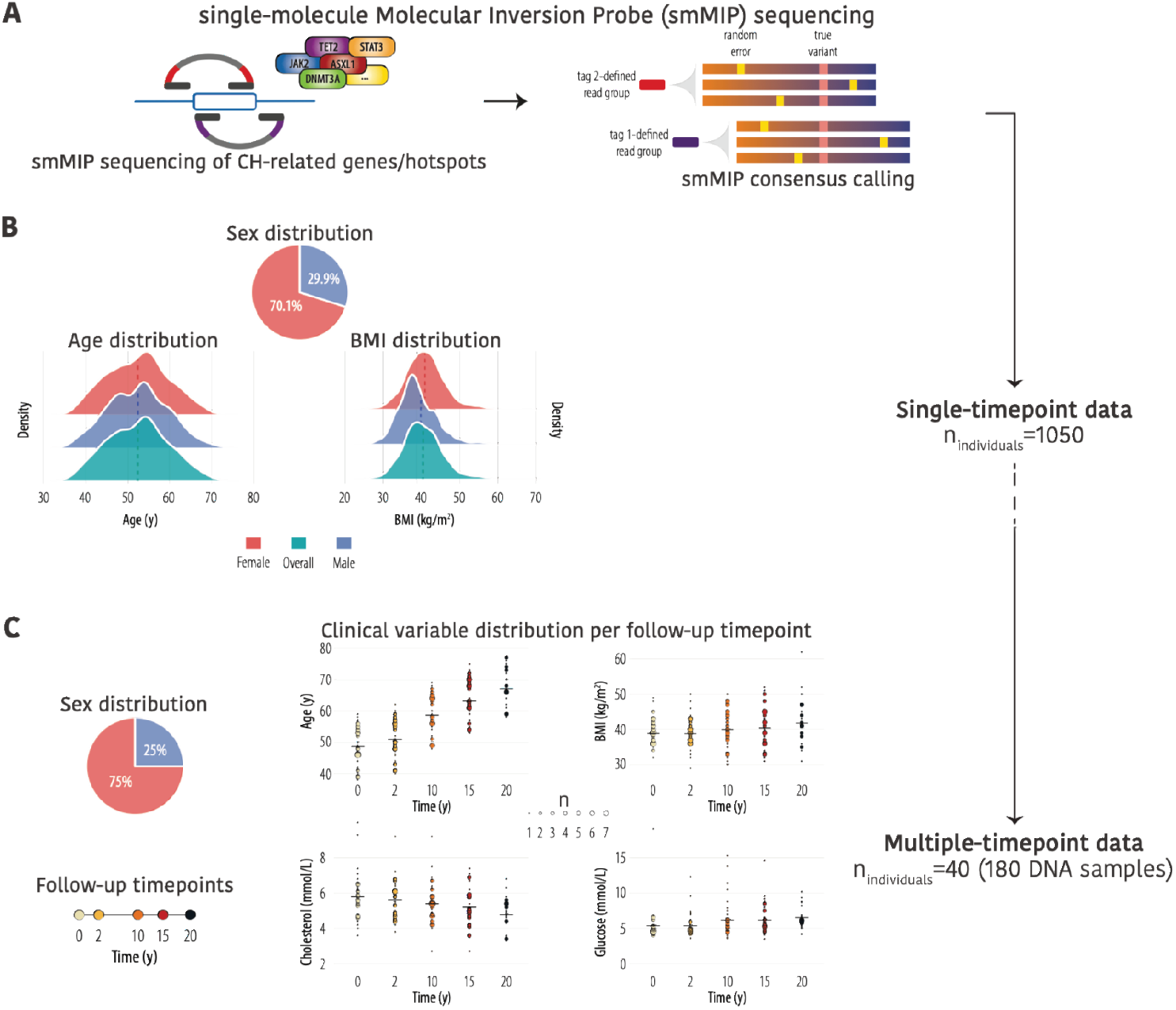
CHDM identification in single- and multiple-timepoint datasets. **(A)** Single- molecule molecular inversion probes (smMIPs) were used for ultra-sensitive and error- corrected targeted sequencing of 24 clonal haematopoiesis-related genes/hotspots to enable reliable detection of CHDMs. For more details, see Supplementary material, File S2. **(B)** Sex, age and BMI distribution in the single-timepoint dataset (1050 individuals). **(C)** Sex distribution and distribution of clinical variables at the indicated follow-up timepoints in the multiple-timepoint dataset (40 individuals).

The age of individuals in our single-timepoint dataset at the time of DNA sampling ranged from 37.3 to 71.2 years (mean 52.4 years). Women were overrepresented (70.1% vs 29.9% men), but the age distribution was comparable in women and men (Figure 1B). Anthropomorphic measurements at baseline reflected the inclusion criteria for this cohort with obesity (Figure 1B and Supplementary material online, *File S3*).

For the 20-year multiple-timepoint dataset with data from 40 individuals, DNA was available for analysis from five timepoints (baseline/0-, 2-, 10-, 15-, 20 years) for more than half of the individuals (n=21), from four timepoints for 18 individuals, and from three timepoints for one individual. Women were again overrepresented (75.0% vs 25.0% men) (Figure 1C). BMI, serum cholesterol, and plasma glucose levels remained relatively stable over 20 years (Figure 1C).

### CHDM prevalence and clone size in the single-timepoint database

In our single-timepoint dataset of 1050 individuals, we identified a total of 273 candidate CHDMs, consisting of 135 different mutations, in 216 individuals (Supplementary material online, *File S3*). A single CHDM was detected in 172 individuals (16.4%), and 44 individuals (4.2%) carried more than one CHDM (Figure 2A).

**Figure 2.**
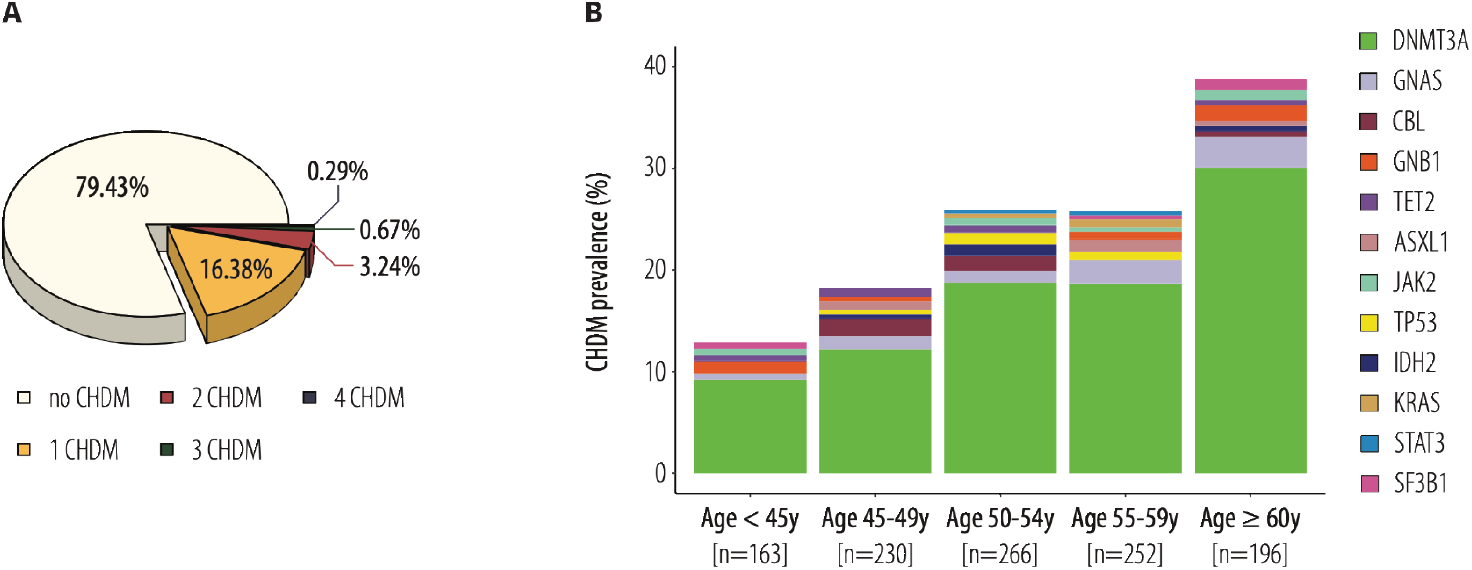
CHDM prevalence and affected genes in the single-timepoint dataset. **(A)** Percentage of individuals (total n = 1050) with no, a single, two, three or four CHDMs. **(B)** Total CHDM prevalence and the affected genes per age category.

The majority of mutations identified in this study have previously been described: 226 (82.8% of the single-timepoint mutations) were identical to previously identified CHDMs from the literature or showed a novel loss-of-function (LoF) mutation in *DNTM3A, TET2* or *ASXL1* (Supplementary material online, *File S3*). For details on identified CHDMs and comparison with the literature see Supplementary material online, *File S5*).

We observed an increasing prevalence of CHDMs with age (Figure 2B), and CHDM carriers were significantly older than non-carriers (54.6 and 51.9 years, respectively, P=2.43E^-06^). Mutations in *DNMT3A* were the most common in all age categories (Figure 2B).

The VAF of all 273 candidate CHDMs ranged from 0.01% to 31.15% (mean 2.72%, median 0.70%); 203 of the CHDMs detected had a VAF <2%, *i*.*e*. below the arbitrary threshold for defining mutations as CHIP, highlighting the sensitivity of our smMIP assay (Figure 3A).

**Figure 3.**
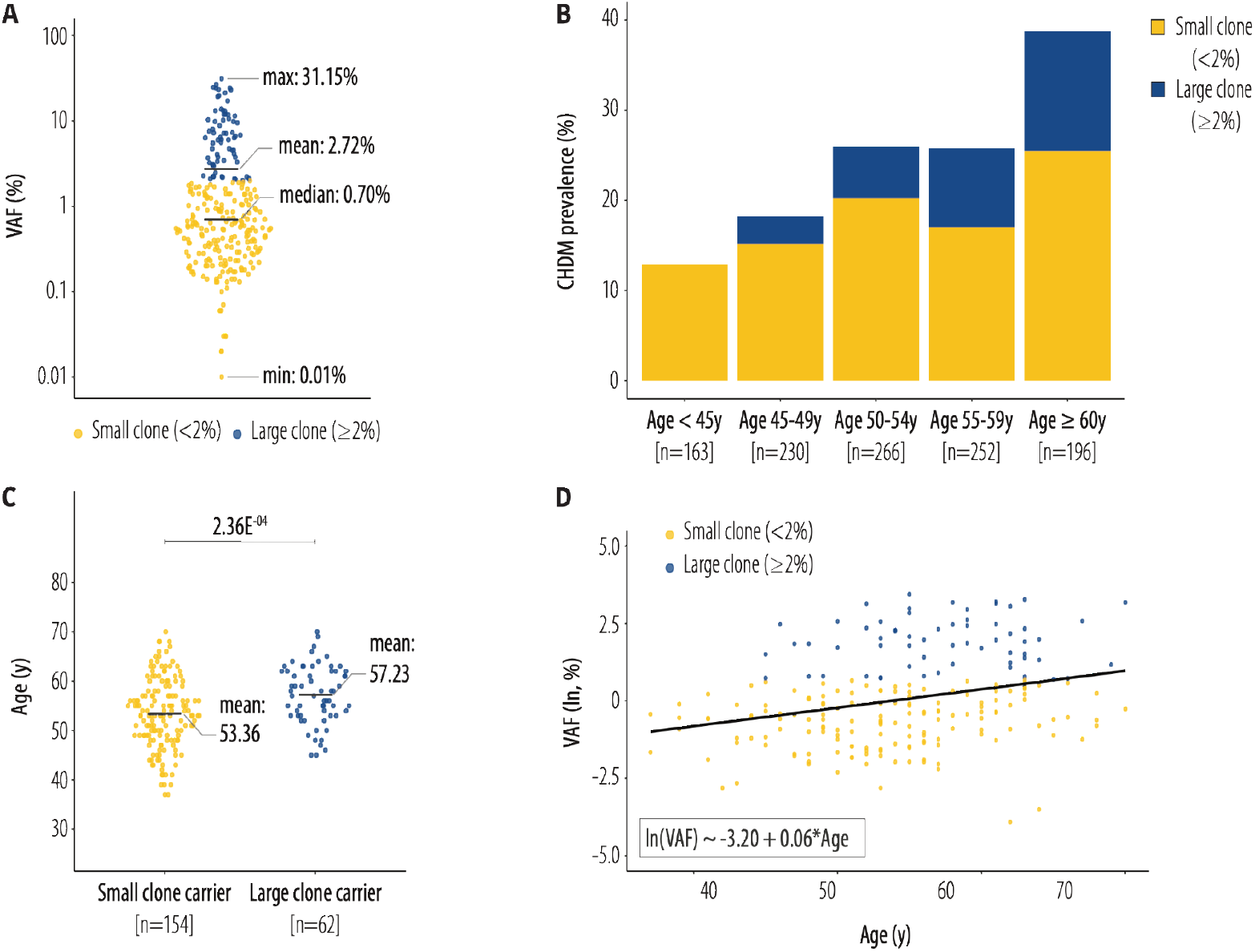
CHDM clone size and relation to age in the single-timepoint dataset. **(A)** The VAF of identified CHDMs. **(B)** Prevalence of small and large clones per age category. **(C)** Age of small-clone carriers and large-clone carriers. **(D)** Visualization of linear regression model using clone size of all CHDMs (log-transformed VAF) as dependent variable and age as predictor. For details on model parameters, see (Supplementary material, File S3).

The prevalence of both small (VAF <2%) and large clones (VAF ≥2%) increased with age, and the proportion of large clones was highest in the oldest age category (Figure 3B).

Accordingly, the mean age was significantly higher in large-versus small-clone carriers (57.2 and 53.4 years, respectively, P=2.36E^-04^) (Figure 3C). The association between clone prevalence and age, as determined by logistic regression models, was stronger when only large clones were considered (odds ratio (OR)_age_=1.05, 95% confidence interval (CI) 1.03-1.08 for all clones; OR_age_=1.11, 95% CI 1.07-1.16 for large clones). By means of linear regression, we also observed a significant association of age with clone size (log-transformed VAF) (Figure 3D) with an effect estimate of 0.059 (P=2.58E^-05^), which translates to a 6% increase in clone size per year.

### Evolution of CHDMs over up to 20 years

A multiple-timepoint dataset was created by selecting 40 individuals in whom we identified at least one CHDM in the single-timepoint dataset and for whom blood samples for DNA extraction were available at three or more timepoints over 20 years. Among these 40 individuals, we identified 115 CHDMs, comprising 53 unique mutations with up to 6 mutations per individual (Supplementary material online, *File S6*). Similar to the single-timepoint results, 71 of the 115 detected mutations (61.7%) have been reported in the literature or represent LoF mutations in *DNMT3A, TET2* or *ASXL1*. Of the 53 unique mutations, 39 (73.58%) have previously been reported or represent novel LoF mutations in these three genes (Supplementary material online, *File S3*).

The CHDMs evolved over time, but this evolution varied substantially both between individuals and mutations (Supplementary material online, *File S6*). Figure 4 shows examples of CHDM evolution over 20 years for three individuals. One individual had six different clones, none of which seemed to have a clear advantage over the others (Figure 4A). In contrast, in another individual, *GNB1* [p.(Lys57Glu)] appeared to have a growth advantage, as *DNMT3A* [p.(Asp702Glu)] started shrinking when *GNB1* [p.(Lys57Glu)] expanded (Figure 4B). In a third individual, we detected four CHDMs, two of which (*TET2* [p.(Glu873^*^)] and *DNMT3A* [p.(Phe732Ile)]) grew to a VAF of almost 40% after 20 years, but were far below the CHIP level at the first time of detection (Figure 4C).

**Figure 4.**
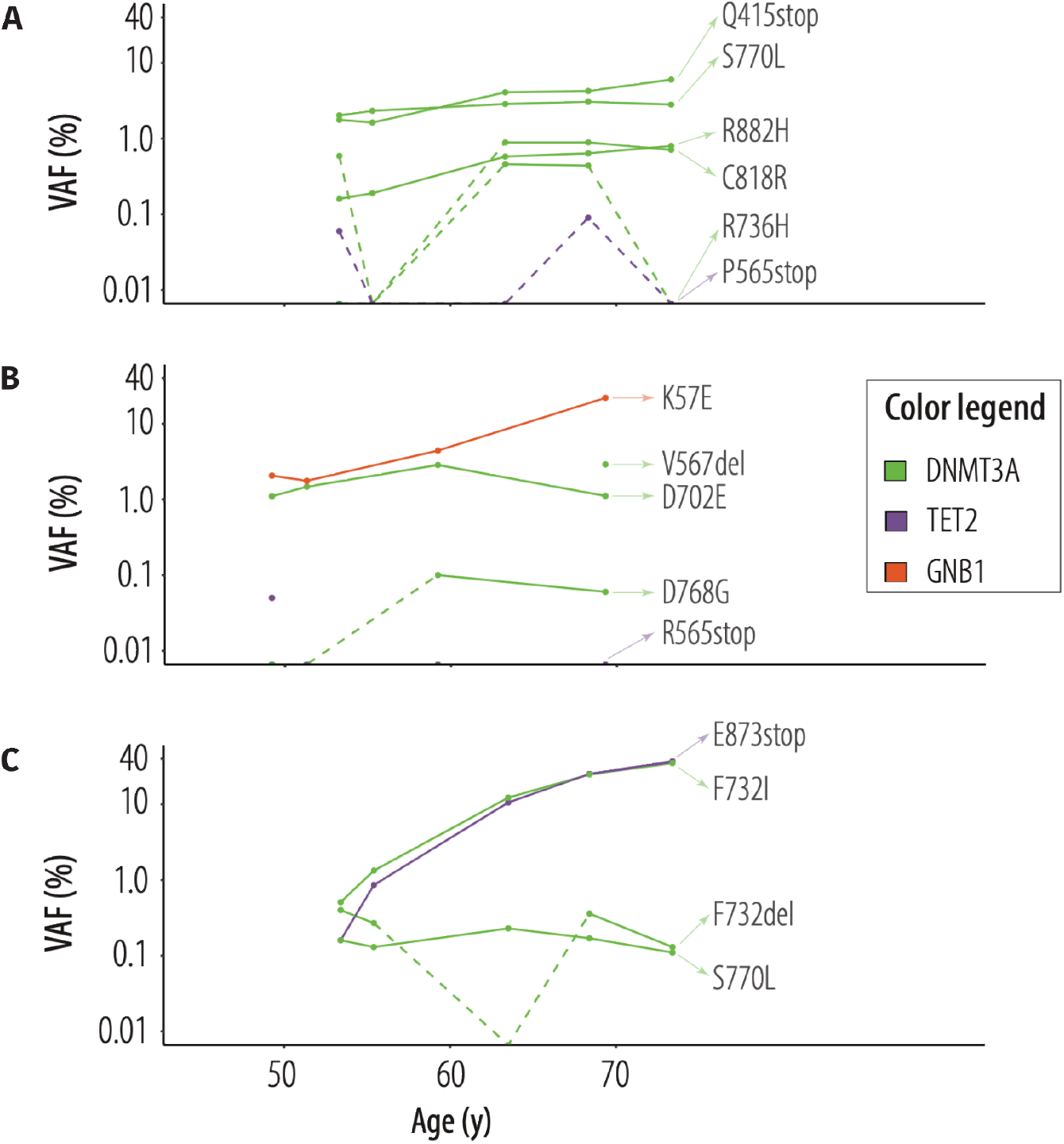
CHDM evolution over time in the multiple-timepoint dataset. Examples of CHDM evolution over 20 years in three individuals with obesity. Solid lines connect mutations detected in consecutive samples while broken lines represent mutations that were undetectable at some time points. **(A)** This individual had five mutations in *DNMT3A* and one in *TET2*, with similar growth patterns over time. **(B)** This individual had two mutations in *DNMT3A*, one in *GNB1*, and one in *TET2*. The clone [p.(Lys57Glu)] in *GNB1* appeared to have a growth advantage over [p.(Asp702Glu)] in *DNMT3A*. **(C)** This individual had two mutations in *TET2* and two mutations in *DNMT3A*. All clones were below the CHIP level when first detected, but two of them, [p.(Glu873*)] in *TET2* and [p.(Phe732Ile)] in *DNMT3A*, grew to a VAF of almost 40% over 20 years.

The mean VAF of all detected CHDMs in the 40 individuals increased with follow-up time (Figure 5A). Of the 115 identified CHDMs, 38 were categorized as events and 77 were defined as traceable trajectories (Figure 5B), with 22 individuals carrying more than one trajectory.

**Figure 5.**
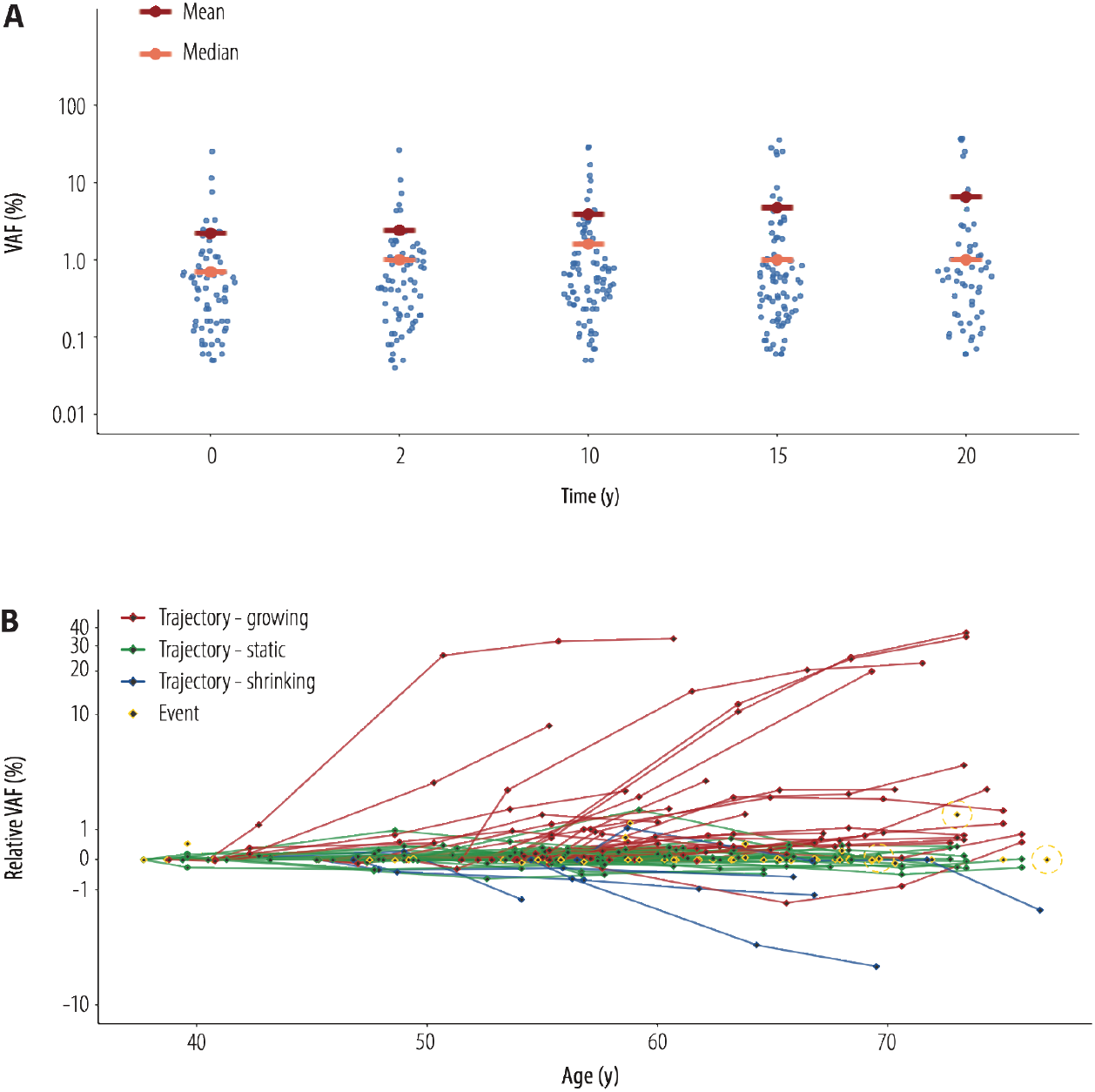
Clone size of CHDM at multiple-timepoints over 20 years. **(A)** The variant allele frequency (VAF) of all detected CHDMs per follow-up timepoint, with mean and median VAF indicated. **(B)** Relative changes in clone size of CHDMs over 20 years and according to chronological age. Trajectories are connected by red, green and blue lines indicating growing, static and shrinking trajectories, respectively. Single- or multiple-timepoint events are shown in yellow, dashed circles indicate late-appearing clones.

By inspecting the dynamics of all 77 trajectories, we identified 30 growing, 42 static, and 5 shrinking trajectories (Figure 6). In addition, three individuals had late-appearing clones – CHDMs that appeared at the last measured timepoint as a large clone (VAF ≥2%).

**Figure 6.**
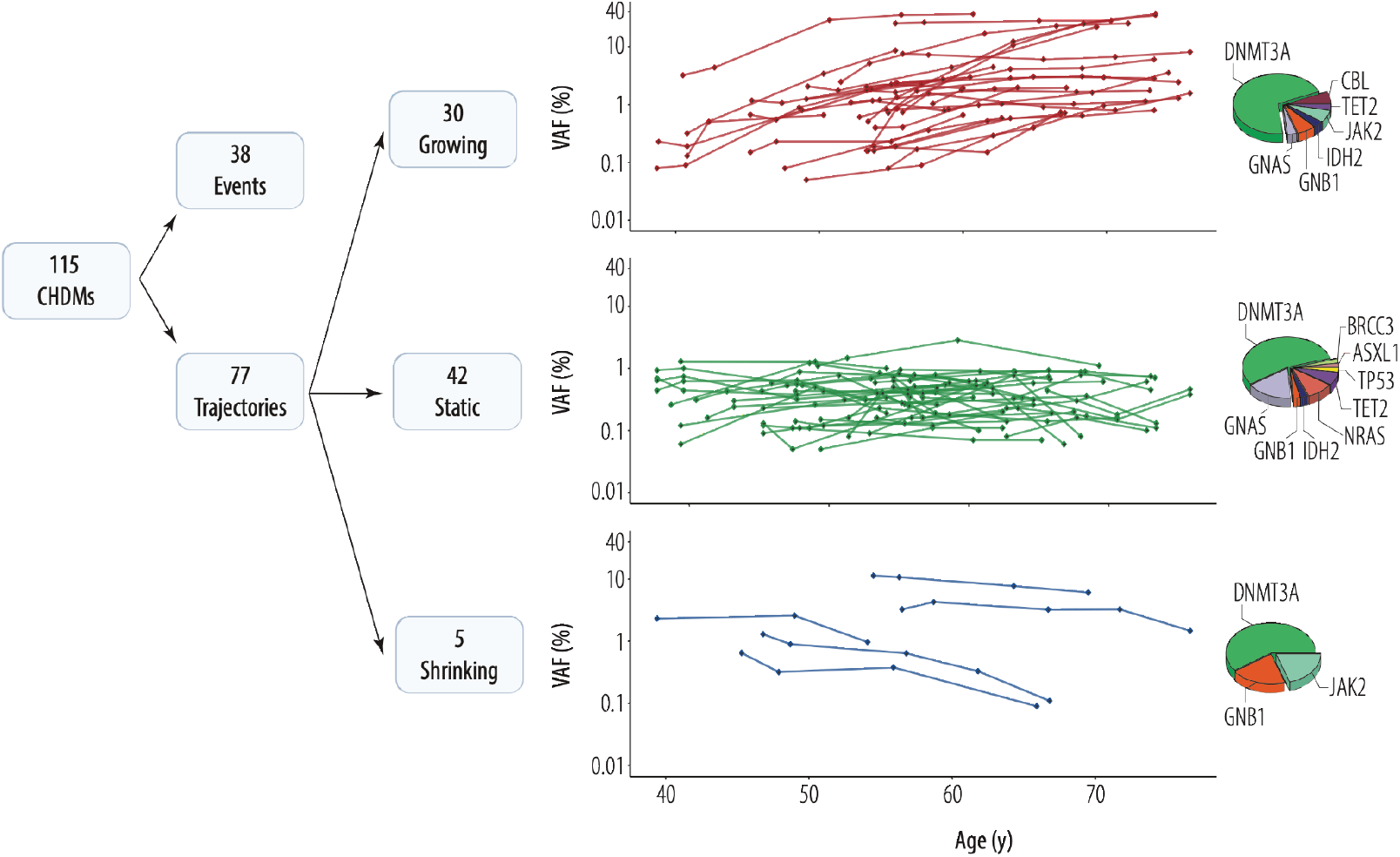
Categorization of CHDMs over 20 years. Categorization of CHDMs into events (single or multiple) and trajectories. Trajectories are further categorized into growing, static and shrinking, and their VAF as well as the proportion of genes in which these trajectories were detected are shown.

The 30 growing trajectories were detected in 21 individuals, carrying mutations in *DNMT3A, CBL, GNAS, JAK2, TET2, GNB1*, and *IDH2*. For 24 growing trajectories, the CHDM was initially detected as a small clone (VAF <2%) and nine of these had become large clones (VAF ≥2%, CHIP) at the last timepoint of detection. For the remaining six growing trajectories, the clone was already above the CHIP level at the first timepoint of detection. The 42 static CHDM trajectories were detected in 24 individuals, carrying mutations in *ASXL1, BRCC3, DNMT3A, GNAS, GNB1, IDH2, NRAS, TET2*, and *TP53*. With the exception of one timepoint in one individual, all static CHDM trajectories were below the CHIP level. The five shrinking CHDM trajectories were detected in five individuals, carrying mutations in *DNMT3A, GNB1*, and *JAK2*. Three started as large clones, two of which shrunk below CHIP.

### Clinical correlations of CHDMs over time

Our multiple-timepoint dataset enabled us to go beyond clone size and examine a potential association between age and actual clone growth. As some individuals carried multiple trajectories over the course of 20 years (Supplementary material online, *File S6*), we first selected the most important trajectory per individual, prioritizing growing clones and clones with largest VAF at any timepoint (Supplementary material online, *File S1*). As we aimed to identify factors associated with clone growth, we excluded individuals in whom the most important trajectory was shrinking (n=5). Because we expected variation in growth patterns in our 32 selected trajectories (Supplementary material online, *File S7*), we used a mixed linear model including random intercept and random slope, to determine the association between age and clonal growth. This model outperformed alternatives with fixed effect estimates as indicated by the Akaike information criterium (Supplementary material online, *File S3*). We identified an average proportionate increase of 7% in VAF per year, ranging from -4% to 27% (Figure 7A) confirming the expected differences in rate of growth per trajectory. These differences in clone growth between individuals were also observed for five identical mutations (Supplementary material online, *File S8*). For example, trajectories of *DNMT3A* [p.(Arg882Cys)], identified in four individuals, and *DNMT3A* [p.(Arg882His)], identified in two individuals, increased with different rates (Figure 7B).

**Figure 7.**
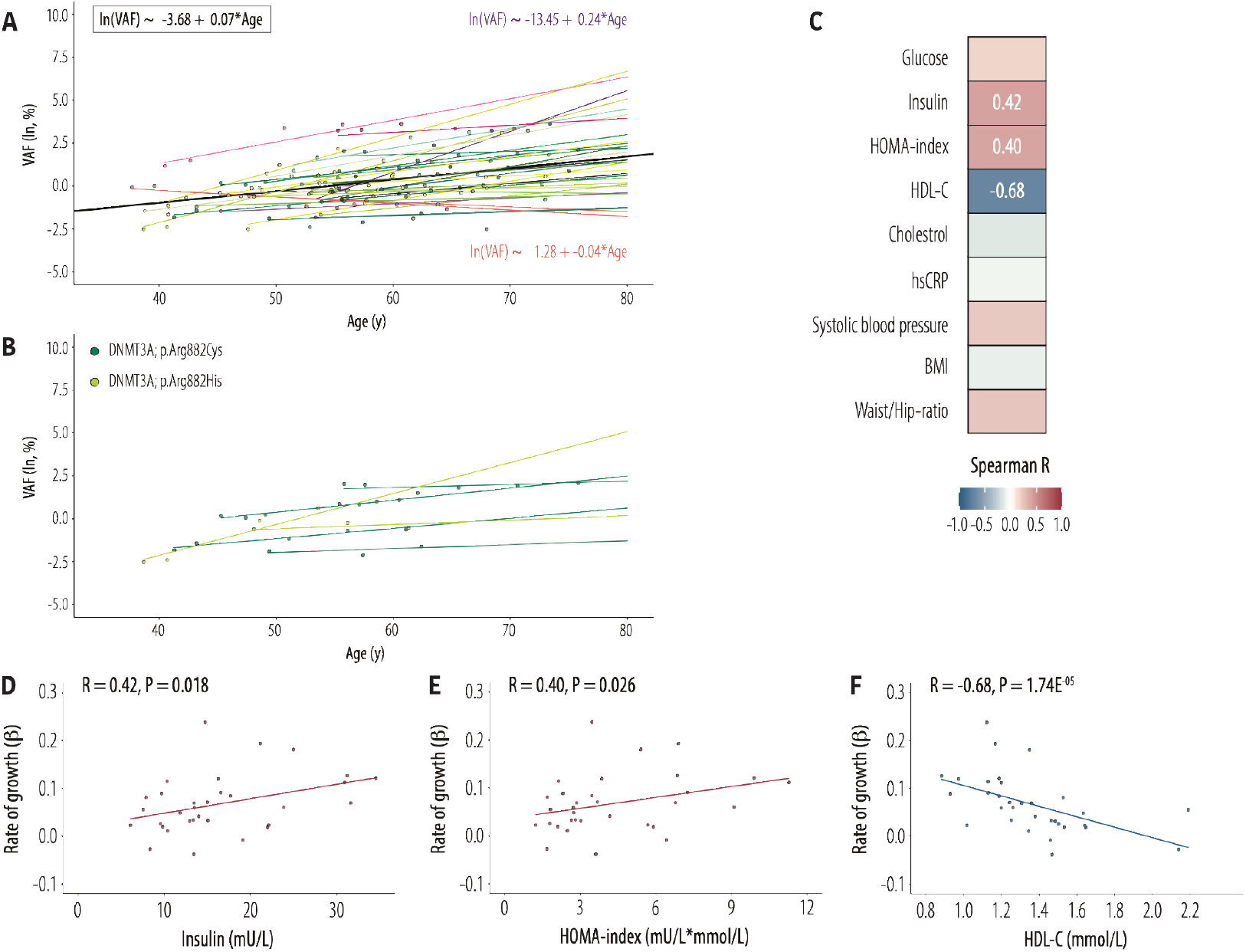
Rate of CHDM growth in relation to age and metabolic dysfunction. **(A)** VAF in relation to age for the 32 most important trajectories. The average mixed linear model (MLM) regression line and equation is shown in black. All individual MLM regression lines are shown in various colors, in purple the trajectory with maximum slope and accompanying equation, in orange the trajectory with minimum slope and accompanying equation. For details on model parameters see Supplementary material, File S3. **(B)** VAFs and slopes for CHDMs in *DNMT3A* [p.(Arg822Cys)] in 4 individuals and *DNMT3A* [p.(Arg882His)] in 2 individuals. **(C)** Heatmap of Spearman R correlations between all individual trajectory effect estimates from our MLM to averaged (based on the first three follow-up timepoints) metabolic clinical parameters. **(D)** Positive Spearman R correlation between individual trajectory effect estimates and insulin. **(E)** Positive Spearman **R** correlation between individual trajectory effect estimates and HOMA-index. **(F)** Negative Spearman R correlation between individual trajectory effect estimates and HDL-C.

We next examined if clinical characteristics were associated with these different rates of growth, as expressed by the effect of age, by correlating all individual trajectory effect estimates from our mixed linear model to metabolic status estimated by cardiovascular risk factors averaged over the first three follow-up timepoints (Figure 7C). We identified significant positive correlations for rate of growth with insulin (Spearman R=0.42, P=0.018) and with insulin resistance measured by HOMA-index (Spearman R=0.40, P=0.025), whereas HDL-C negatively correlated to the rate of growth (Spearman R=-0.68, P=1.74E^-05^) (Figure 7D-F).

## DISCUSSION

In this study we generated and analysed single- and longitudinal multiple-timepoint data on CHDMs in middle-aged individuals with obesity. Approximately 20% of individuals from our cohort had CHDMs detectable by an ultrasensitive assay designed to target known and novel candidate driver mutations. Both CHDM prevalence and clone size increased with age. Importantly, a significant fraction of clones grew, including several initially small clones. In addition to the age-related increase in clone size, clone growth was further accelerated in individuals with metabolic dysfunction.

For the CHDMs identified in the single-timepoint dataset, the clone size was on average 6% larger for each year of age. However, a unique feature of our study was the collection of up to five samples taken over 20 years from 40 well-characterized individuals, which allowed measurement of clonal evolution over time and determination of actual clone growth. When we assessed growth of specific clones within individuals, the average annual growth was 7%. However, there were large inter-individual differences in the rate of clone growth, even between individuals sharing the same CHDM, suggesting that factors other than the mutation itself influence clone growth.

More than half of the individuals in the multiple-timepoint dataset, who were selected on the basis that clones were present, had growing clones, which is an unexpectedly high proportion^18, 20, 21^. This could be explained by the long timeframe of tracing and/or by the severe obesity and obesity-related metabolic abnormalities in our cohort, in contrast to previous longitudinal studies of mainly healthy individuals. Importantly, nine out of the 30 growing trajectories identified in our study were initially small clones that grew over the sampling period to beyond the CHIP level. Our longitudinal data thus showed that clones can be detected many years before they reach the CHIP level.

Our longitudinal observations indicated that increased rate of clone growth was associated not only with age but also with insulin resistance and low HDL-C, both well-established risk factors for cardiovascular disease^22^. These findings are of interest in light of the recent observation that humans with atherosclerosis have increased HSC division rates^23^. Increased HSC proliferation may promote clonal haematopoiesis both by increasing the risk of acquiring CHDMs and by facilitating the expansion of mutant clones^23^. Animal studies indicate that HDL and the adenosine triphosphate–binding cassette (ABC) cholesterol efflux transporter ABCG1, which promotes cholesterol efflux from macrophages to HDL, inhibit HSC proliferation and thereby reduce circulating numbers of leukocytes^24^. A recent Mendelian randomization study in humans supports a causal inverse relationship between HDL-C and leukocyte counts^25^. Reduced HDL efflux capacity is likely to impact atherosclerosis by several mechanisms including both direct effects on foam cells and inflammation in the atherosclerotic plaque but also through enhanced myelopoiesis and platelet production^26^, and this risk may be further increased by CHIP. Similarly, insulin resistance has been associated with high white blood cell count in humans^27-29^. It is therefore possible that the observed association between metabolic dysfunction and clone growth is linked to increased HSC proliferation by directly influencing function and stability of CHIP-related proteins, including TET2^30, 31^. A combination of our results and earlier studies showing that loss of *Tet2* function in HSCs leads to increased atherosclerosis^7, 8^ and insulin resistance^9^ suggest that a vicious cycle might exist, whereby dysfunctional metabolism increases the risk of developing pro-inflammatory CHIP that, once formed, further increases the risk of worsening insulin resistance and atherosclerosis.

The importance of clonal haematopoiesis has mainly been studied for mutations above the CHIP level of 2% because of: (1) methodological difficulties in detecting small clones and (2) the assumption that clones must reach a critical size to be clinically meaningful. CHIP mutations are associated with a wide range of diseases, including haematological malignancies and cardiovascular disease^16^. However, the health consequences of smaller clones are largely unknown and it is unclear how mutations with low VAF evolve over time. Our ultrasensitive assay allowed us to detect small clones, and almost 75% of all CHDMs in the single-timepoint data were below the CHIP level definition. Irrespective of clone size, approximately 80% of all clones that we found overlapped with CHDMs that have been described in the literature, indicating that the higher sensitivity of our assay did not result in increased false positive findings. In addition, we observed an age-related increase in the prevalence of both small and large clones, and repeated measurements over two decades showed that small clones may grow beyond the CHIP level. This indicates that the presence of small clones is also linked to ageing and possibly age-related phenotypes, and that with time small clones may become large enough to become clinically relevant. The importance of small clones is supported by a recent study in which VAFs significantly lower than the CHIP level for *DNMT3A* and *TET2* mutations were associated with impaired survival in patients with chronic ischemic heart failure^17^.

CHDM prevalence comparisons are heavily dependent on the technology applied^32^. The ultra-sensitive assay used in this study detects smaller clones as well as additional mutations in the full coding sequence of *DNMT3*. Our previous work using an assay similar to that used in the current study indicated an overall prevalence of CHDMs of 9.6% in 2,000 individuals from the general Dutch population with an average age of 45 years, while a subgroup of 400 individuals with age similar to the current cohort had a prevalence of 13.2%^19^. The 20% CHDM prevalence in the single-timepoint dataset, with an average age of 53 years, is high compared to most other reports^32, 33^. This is consistent with higher clonal haematopoiesis in individuals with obesity, in line with one recent study^15^. Furthermore, we observed mean VAF increases of 6-7% per year, which are higher than earlier reports of 2-4% annual increase in healthy individuals^23^ and again support potentially greater clone growth in individuals with obesity. Obesity is a complex phenotype, often associated with a wide range of metabolic abnormalities and diseases, accelerated aging and shortened lifespan. Accumulating evidence shows that there must be factors beyond traditional risk factors that contribute to the link between obesity and cardiovascular disease^1, 33^. It is therefore tempting to speculate that clonal haematopoiesis could in part explain increased cardiovascular disease and decreased life expectancy in people with obesity.

The increased risk associated with CHIP for several severe diseases and overall mortality raises the question of how this risk could be mitigated. In individuals with established CHIP, interventions may target the pro-inflammatory activity of CHIP^2, 34^. Mice with experimental CHIP show increased pro-inflammatory drive of the NLRP3/interleukin (IL)-1 /IL-6 axis and atherosclerosis^33, 35^, and humans carrying a genetic deficiency in *IL6* appear to have reduced cardiovascular risk related to CHIP^36^. Thus, intervention with IL-6 inhibitors could potentially reduce the risk of cardiovascular disease in patients with CHIP. In addition, individualized treatments that increase HDL-C and/or insulin sensitivity may potentially target the risk associated with CHIP by preventing expansion of small clones or even reducing the size of clones in patients with CHIP.

Limitations of this study include the use of a targeted assay which may miss mutations in other genes or outside targeted hotspots. Furthermore, our study is limited by the lack of individuals with normal body weight for comparison. Strengths of this study are the ability to detect small clones down to VAF 0.01%, and the longitudinal measurements in general, which allowed us to examine how CHDMs develop over up to 20 years and to examine the association with clinical characteristics.

In conclusion, this study showed that approximately one in five middle-aged individuals with obesity in our cohort had detectable CHDMs, and that a proportion of these clones expanded to become CHIP, mutations that have been linked to haematological cancers and cardiovascular and metabolic diseases. We also demonstrated that accelerated growth of clones was associated with insulin resistance and low HDL-C, suggesting biological mechanisms that may open up opportunities for strategies to reduce the risk of clones expanding to CHIP level.

## Supporting information

Supplementary materials and methods, results, references

## Acknowledgments

We thank the Radboud Genomics Technology Center, Radboud University Medical Center Nijmegen, for their technical assistance.

## Data availability

Code for processing and filtering smMIP-based sequencing data is extensively explained in the methods section and supplementary information of this manuscript and will be made available upon reasonable request. The source code from the R-packages that were used in this study are freely available online (https://cran.r-project.org/).

The data are subject to legal restrictions according to national legislation. Confidentiality regarding personal information in studies is regulated in the Public Access to Information and Secrecy Act (SFS 2009:400), OSL. A request to get access to public documents can be rejected or granted with reservations by the University of Gothenburg. If University of Gothenburg refuses to disclose the documents the applicant is entitled to get a written decision that can be appealed to the administrative court of appeal.

## References

1. Fuster JJ, Walsh K. Somatic Mutations and Clonal Hematopoiesis: Unexpected Potential New Drivers of Age-Related Cardiovascular Disease. Circ Res 2018;122(3):523–532.

2. Jaiswal S, Libby P. Clonal haematopoiesis: connecting ageing and inflammation in cardiovascular disease. Nat Rev Cardiol 2020;17(3):137–144.

3. Jaiswal S, Fontanillas P, Flannick J, Manning A, Grauman PV, Mar BG, Lindsley RC, Mermel CH, Burtt N, Chavez A, Higgins JM, Moltchanov V, Kuo FC, Kluk MJ, Henderson B, Kinnunen L, Koistinen HA, Ladenvall C, Getz G, Correa A, Banahan BF, Gabriel S, Kathiresan S, Stringham HM, McCarthy MI, Boehnke M, Tuomilehto J, Haiman C, Groop L, Atzmon G, Wilson JG, Neuberg D, Altshuler D, Ebert BL. Age-related clonal hematopoiesis associated with adverse outcomes. N Engl J Med 2014;371(26):2488–98.

4. Wong TN, Miller CA, Jotte MRM, Bagegni N, Baty JD, Schmidt AP, Cashen AF, Duncavage EJ, Helton NM, Fiala M, Fulton RS, Heath SE, Janke M, Luber K, Westervelt P, Vij R, DiPersio JF, Welch JS, Graubert TA, Walter MJ, Ley TJ, Link DC. Cellular stressors contribute to the expansion of hematopoietic clones of varying leukemic potential. Nat Commun 2018;9(1):455.

5. Steensma DP, Bejar R, Jaiswal S, Lindsley RC, Sekeres MA, Hasserjian RP, Ebert BL. Clonal hematopoiesis of indeterminate potential and its distinction from myelodysplastic syndromes. Blood 2015;126(1):9–16.

6. Bonnefond A, Skrobek B, Lobbens S, Eury E, Thuillier D, Cauchi S, Lantieri O, Balkau B, Riboli E, Marre M, Charpentier G, Yengo L, Froguel P. Association between large detectable clonal mosaicism and type 2 diabetes with vascular complications. Nat Genet 2013;45(9):1040–3.

7. Jaiswal S, Natarajan P, Silver AJ, Gibson CJ, Bick AG, Shvartz E, McConkey M, Gupta N, Gabriel S, Ardissino D, Baber U, Mehran R, Fuster V, Danesh J, Frossard P, Saleheen D, Melander O, Sukhova GK, Neuberg D, Libby P, Kathiresan S, Ebert BL. Clonal Hematopoiesis and Risk of Atherosclerotic Cardiovascular Disease. N Engl J Med 2017;377(2):111–121.

8. Wang Y, Sano S, Yura Y, Ke Z, Sano M, Oshima K, Ogawa H, Horitani K, Min KD, Miura-Yura E, Kour A, Evans MA, Zuriaga MA, Hirschi KK, Fuster JJ, Pietras EM, Walsh K. Tet2-mediated clonal hematopoiesis in nonconditioned mice accelerates age-associated cardiac dysfunction. JCI Insight 2020;5(6).

9. Fuster JJ, Zuriaga MA, Zorita V, MacLauchlan S, Polackal MN, Viana-Huete V, Ferrer-Perez A, Matesanz N, Herrero-Cervera A, Sano S, Cooper MA, Gonzalez-Navarro H, Walsh K. TET2-Loss-of-Function-Driven Clonal Hematopoiesis Exacerbates Experimental Insulin Resistance in Aging and Obesity. Cell Rep 2020;33(4):108326.

10. Tzanetakou IP, Katsilambros NL, Benetos A, Mikhailidis DP, Perrea DN. “Is obesity linked to aging?”: adipose tissue and the role of telomeres. Ageing Res Rev 2012;11(2):220–9.

11. Bhaskaran K, Dos-Santos-Silva I, Leon DA, Douglas IJ, Smeeth L. Association of BMI with overall and cause-specific mortality: a population-based cohort study of 3.6 million adults in the UK. Lancet Diabetes Endocrinol 2018;6(12):944–953.

12. Carlsson LMS, Sjoholm K, Jacobson P, Andersson-Assarsson JC, Svensson PA, Taube M, Carlsson B, Peltonen M. Life Expectancy after Bariatric Surgery in the Swedish Obese Subjects Study. N Engl J Med 2020;383(16):1535–1543.

13. Prospective Studies C, Whitlock G, Lewington S, Sherliker P, Clarke R, Emberson J, Halsey J, Qizilbash N, Collins R, Peto R. Body-mass index and cause-specific mortality in 900 000 adults: collaborative analyses of 57 prospective studies. Lancet 2009;373(9669):1083–96.

14. Valdes AM, Andrew T, Gardner JP, Kimura M, Oelsner E, Cherkas LF, Aviv A, Spector TD. Obesity, cigarette smoking, and telomere length in women. Lancet 2005;366(9486):662–4.

15. Haring B, Reiner AP, Liu J, Tobias DK, Whitsel E, Berger JS, Desai P, Wassertheil-Smoller S, LaMonte MJ, Hayden KM, Bick AG, Natarajan P, Weinstock JS, Nguyen PK, Stefanick M, Simon MS, Eaton CB, Kooperberg C, Manson JE. Healthy Lifestyle and Clonal Hematopoiesis of Indeterminate Potential: Results From the Women’s Health Initiative. J Am Heart Assoc 2021;10(5):e018789.

16. Silver AJ, Jaiswal S. Clonal hematopoiesis: Pre-cancer PLUS. Adv Cancer Res 2019;141:85–128.

17. Assmus B, Cremer S, Kirschbaum K, Culmann D, Kiefer K, Dorsheimer L, Rasper T, Abou-El-Ardat K, Herrmann E, Berkowitsch A, Hoffmann J, Seeger F, Mas-Peiro S, Rieger MA, Dimmeler S, Zeiher AM. Clonal haematopoiesis in chronic ischaemic heart failure: prognostic role of clone size for DNMT3A- and TET2-driver gene mutations. Eur Heart J 2021;42(3):257–265.

18. Young AL, Challen GA, Birmann BM, Druley TE. Clonal haematopoiesis harbouring AML-associated mutations is ubiquitous in healthy adults. Nat Commun 2016;7:12484.

19. Acuna-Hidalgo R, Sengul H, Steehouwer M, van de Vorst M, Vermeulen SH, Kiemeney L, Veltman JA, Gilissen C, Hoischen A. Ultra-sensitive Sequencing Identifies High Prevalence of Clonal Hematopoiesis-Associated Mutations throughout Adult Life. Am J Hum Genet 2017;101(1):50–64.

20. van den Akker EB, Makrodimitris S, Hulsman M, Brugman MH, Nikolic T, Bradley T, Waisfisz Q, Baas F, Jakobs ME, de Jong D, Slagboom PE, Staal FJT, Reinders MJT, Holstege H. Dynamic clonal hematopoiesis and functional T-cell immunity in a supercentenarian. Leukemia 2020.

21. Young AL, Tong RS, Birmann BM, Druley TE. Clonal hematopoiesis and risk of acute myeloid leukemia. Haematologica 2019;104(12):2410–2417.

22. Kopin L, Lowenstein C. Dyslipidemia. Ann Intern Med 2017;167(11):ITC81–ITC96.

23. Heyde A, Rohde D, McAlpine CS, Zhang S, Hoyer FF, Gerold JM, Cheek D, Iwamoto Y, Schloss MJ, Vandoorne K, Iborra-Egea O, Munoz-Guijosa C, Bayes-Genis A, Reiter JG, Craig M, Swirski FK, Nahrendorf M, Nowak MA, Naxerova K. Increased stem cell proliferation in atherosclerosis accelerates clonal hematopoiesis. Cell 2021;184(5):1348–1361 e22.

24. Yvan-Charvet L, Pagler T, Gautier EL, Avagyan S, Siry RL, Han S, Welch CL, Wang N, Randolph GJ, Snoeck HW, Tall AR. ATP-binding cassette transporters and HDL suppress hematopoietic stem cell proliferation. Science 2010;328(5986):1689–93.

25. Harslof M, Pedersen KM, Nordestgaard BG, Afzal S. Low HDL (High-Density Lipoprotein) Cholesterol and High White Blood Cell Counts: A Mendelian Randomization Study. Arterioscler Thromb Vasc Biol 2020:ATVBAHA120314983.

26. Murphy AJ, Tall AR. Disordered haematopoiesis and athero-thrombosis. Eur Heart J 2016;37(14):1113–21.

27. Facchini F, Hollenbeck CB, Chen YN, Chen YD, Reaven GM. Demonstration of a relationship between white blood cell count, insulin resistance, and several risk factors for coronary heart disease in women. J Intern Med 1992;232(3):267–72.

28. Kuo TY, Wu CZ, Lu CH, Lin JD, Liang YJ, Hsieh CH, Pei D, Chen YL. Relationships between white blood cell count and insulin resistance, glucose effectiveness, and first- and second-phase insulin secretion in young adults. Medicine (Baltimore) 2020;99(43):e22215.

29. Piedrola G, Novo E, Escobar F, Garcia-Robles R. White blood cell count and insulin resistance in patients with coronary artery disease. Ann Endocrinol (Paris) 2001;62(1 Pt 1):7–10.

30. Lee MKS, Dragoljevic D, Bertuzzo Veiga C, Wang N, Yvan-Charvet L, Murphy AJ. Interplay between Clonal Hematopoiesis of Indeterminate Potential and Metabolism. Trends Endocrinol Metab 2020;31(7):525–535.

31. Wu D, Hu D, Chen H, Shi G, Fetahu IS, Wu F, Rabidou K, Fang R, Tan L, Xu S, Liu H, Argueta C, Zhang L, Mao F, Yan G, Chen J, Dong Z, Lv R, Xu Y, Wang M, Ye Y, Zhang S, Duquette D, Geng S, Yin C, Lian CG, Murphy GF, Adler GK, Garg R, Lynch L, Yang P, Li Y, Lan F, Fan J, Shi Y, Shi YG. Glucose-regulated phosphorylation of TET2 by AMPK reveals a pathway linking diabetes to cancer. Nature 2018;559(7715):637–641.

32. Watson CJ, Papula AL, Poon GYP, Wong WH, Young AL, Druley TE, Fisher DS, Blundell JR. The evolutionary dynamics and fitness landscape of clonal hematopoiesis. Science 2020;367(6485):1449–1454.

33. Jaiswal S, Ebert BL. Clonal hematopoiesis in human aging and disease. Science 2019;366(6465).

34. Abplanalp WT, Cremer S, John D, Hoffmann J, Schuhmacher B, Merten M, Rieger MA, Vasa-Nicotera M, Zeiher AM, Dimmeler S. Clonal Hematopoiesis-Driver DNMT3A Mutations Alter Immune Cells in Heart Failure. Circ Res 2021;128(2):216–228.

35. Fuster JJ, MacLauchlan S, Zuriaga MA, Polackal MN, Ostriker AC, Chakraborty R, Wu CL, Sano S, Muralidharan S, Rius C, Vuong J, Jacob S, Muralidhar V, Robertson AA, Cooper MA, Andres V, Hirschi KK, Martin KA, Walsh K. Clonal hematopoiesis associated with TET2 deficiency accelerates atherosclerosis development in mice. Science 2017;355(6327):842–847.

36. Bick AG, Pirruccello JP, Griffin GK, Gupta N, Gabriel S, Saleheen D, Libby P, Kathiresan S, Natarajan P. Genetic Interleukin 6 Signaling Deficiency Attenuates Cardiovascular Risk in Clonal Hematopoiesis. Circulation 2020;141(2):124–131.

