## Supplementary materials and methods, results, references for "Expansion of mutation-driven haematopoietic clones is associated with insulin resistance and low HDL-cholesterol in individuals with obesity"

### Supplementary material

| Content | Description |
|---|---|
| **File S1:**<br>Materials and Methods, Results, References | Additional information on the study cohort, data generation and analysis, and additional detailed results.<br>References for the Supplementary material |
| **File S2:**<br>Figure illustrating CHDM detection | smMIP workflow to optimize sensitivity for low-level somatic mutation detection<br>a: Laboratory procedure<br>b: Mapping and Variant Calling<br>c: Quality Filtering |
| **File S3:**<br>Tables 1-7 | -table 1a: Sequenced gene regions<br>-table 1b: smMIP probes<br>-table 2: Baseline characteristics in the single-timepoint dataset<br>-table 3a: CHDMs in literature<br>-table 3b: Overlap of CHDMs in single-timepoint dataset and CHDMs in literature (shown in table 3a)<br>-table 3c Overlap of CHDMs in multiple-timepoint dataset and CHDMs in literature (shown in table 3a)<br>-table 4: CHDMs identified in the single-timepoint dataset<br>-table 5a-c: Regression model parameters<br>-table 6: CHDMs identified in the multiple-timepoint dataset<br>-table 7a: Mixed Linear Model (MLM) fixed and random parameters<br>-table 7b: Mixed Linear Model comparisons |
| **File S4:**<br>Figure illustrating smMIP coverage | smMIP coverage of single- and multiple-timepoint datasets<br>a: Single-timepoint dataset<br>b: Multiple-timepoint dataset |
| **File S5:**<br>Figure illustrating identified CHDMs and comparison to literature | CHDMs identified in the single- and multiple-timepoint datasets and comparison to literature<br>a: Gene distribution of CHDMs in the single-timepoint dataset<br>b: Gene distribution of CHDMs in the multiple-timepoint dataset<br>c: CHDMs resulting in specific amino acid changes in the single-timepoint dataset<br>d: Amino acid changes caused by *DNMT3A* mutations in the single-timepoint dataset<br>e: Amino acid changes caused by *DNMT3A* mutations in the multiple-timepoint dataset<br>f: DNMT3A mutations observed at least five times in literature annotated on a schematic illustration of the DNMT3A protein |
| **File S6:**<br>Figure illustrating CHDMs over time | CHDMs evolvement during 20 years follow-up |
| **File S7:** | Selected trajectories in multiple-timepoint dataset |

| | |
|---|---|
| Figure illustrating variation in growth patterns of CHDMs | |
| **File S8:**<br>Figure illustrating inter-individual differences in clone growth | Differences in rate of growth for recurrent CHDM trajectories in different individuals<br>a: DMNT3A;p.Arg882His<br>b: DNMT3A;p.Arg882Cys<br>c: DNMT3A;p.Tyr735Cys<br>d: GNB1;p.Lys57Glu<br>e: NRAS;p.Ile21Val |

***Material and Methods:***

**Subjects and samples**

*Study cohort*

The SOS study is an ongoing, prospective, controlled intervention study designed to compare outcomes in patients with obesity treated by bariatric surgery (n=2010) and a matched control group given usual care (n=2037)[1]. Inclusion criteria were age between 37 and 60 years and body-mass index (BMI) of ≥34 for men and ≥38 for women. The exclusion criteria were earlier surgery for gastric or duodenal ulcer, earlier bariatric surgery, gastric ulcer during the past 6 months, ongoing malignancy, active malignancy during the past 5 years, myocardial infarction during the past 6 months, bulimic eating pattern, drug or alcohol abuse, psychiatric or cooperative problems contraindicating bariatric surgery, other contraindicating conditions (such as chronic glucocorticoid or anti-inflammatory treatment). The current study includes individuals from the control group given usual care, with follow-up visits at 480 primary health care centers at baseline (0), 0.5, 1, 2, 3, 4, 6, 8, 10, 15, and 20 years.

**CHDM detection by single-molecule Molecular Inversion probe (smMIP) sequencing**

*DNA extraction*

For most individuals in the single-timepoint dataset a blood sample specifically for genetic analyses was collected between 1998 and 2006. The blood was collected in EDTA-tubes, transferred to plastic bottles and stored in -80°C until DNA extraction. DNA was extracted using the AGOWA mag kit (LGC Group, Teddington, Middlesex, UK). Extractions were performed by the Genomics core facility at Sahlgrenska Academy, University of Gothenburg (Gothenburg, Sweden).

In the multiple-timepoint dataset, DNA was extracted from blood samples routinely collected at the follow-up visits at baseline, 2-, 10-, 15-, and 20-years. Blood was collected in Heparin-tubes, transferred to cryotubes and stored at -80°C until DNA extraction. Extractions were performed by the Qiagen genomics services (Hilden, Germany) using the DNeasy 96 Blood & Tissue kit (Qiagen, Hilden, Germany).

*Sequencing*

CHDMs were analyzed by ultra-sensitive sequencing, as essentially previously described[2]. We modified the existing assay by removing non-CH-related hotspot targets and designing additional double tiling smMIP probes[3] for the entire coding sequence of the most prevalently reported CH-driver gene *DNMT3A*. The final assay consisted of a total of 300 smMIP probes, with 54 nucleotide target sequence per probe, spanning a total of 7612 bases of target sequence (see *File S3, table 1a* for a list of genes, bases and target hotspots covered, and *File S3, table 1b* for a list of all smMIP probes). smMIP captures were likewise performed with slight modifications: 200-300ng gDNA of each sample with a DNA to smMIP ratio of 1:2,400; all samples were captured twice and each replicate was tagged with an independent barcode by PCR. Sequencing was performed in batches of up to 380 samples per run by 2x79 basepair PE reads on a high-output run on a NextSeq500 instrument (Illumina) (*File S2, fig. a*).

*Variant calling*

For the purpose of providing true positive somatic variant calls, we applied two independent data processing strategies followed up by targeted quality control, specifically designed for this study (*File S2, fig. b*). Specifically, FASTQ files were: 1) aligned to the entire reference genome (Hg19) with BWA-MEM[4], and 2) imported into the commercially available NGS software package Sequence Pilot (JSI Medical Systems), using the optimized smMIP analysis module as described previously[5, 6]. The latter allows for a consensus calling per smMIP probe, enhancing individual variant quality by reducing random PCR or sequencing artefacts, using a majority vote of Unique Molecular Identifier (UMI) duplicates. This also enabled the same molecule to be read with forward and reverse sequencing reads, due to 2x79 basepair reads and a 54 nucleotide insert size of gDNA. Variant calling with Sequence Pilot was performed with the following settings: Minimum combined forward and reverse coverage was set to 10 reads, mutation calling required at least 5 consensus reads (forward and reverse reads considered separately) without a minimal % of variant reads, enabling some somatic calls down to 0.01% (depending on locus specific coverage); consensus calling was done with a minimum of 2 consensus UMI reads and by ignoring consensus read threshold of <30% as ‘likely artefacts'; and UMI-tags with “N" bases or low quality were ignored.

*Quality control single-timepoint dataset*

The resulting variant calls were then subjected to the following stringent quality filtering steps (*File S2, fig. c*): First, individuals with an average coverage below 500x based on the untargeted aligned BAM files were excluded (STEP1). Second, only variants called in both technical replicates were kept (STEP2). Third, the remaining duplicate variant calls were further filtered by excluding non-coding, synonymous, and likely germline (variant allele frequency (VAF) ≥40%) variant calls (STEP3). Fourth, variants called in >5% of the individuals that are considered likely run-specific artefacts (excluding most common known drivers) and common smMIP-run artefacts (based on previously processed smMIP-data) were excluded (STEP4). Fifth, remaining variant calls were flagged based on the following characteristics; a) *PTPN11* variants were excluded, due to mapping issues related to homology with various regions in the genome, b) variants with unspecified alternative allele by JSI (N-allele) were excluded, c) variants called in four or more samples with an alternative allele count below 16 when considering forward and reverse reads separately (based on JSI parameters) were excluded, and d) variants called in less than four samples with an alternative allele count below 24 were excluded, and e) based on visual inspection combined with previous validations[2] we flagged likely true positive and likely false positive variants in green and red respectively, excluding all variants with a red flag, overruling any of the previously described flags. Finally, the percentage of alternative alleles for the remaining variant calls was generated using samtools mpileup[7] on the untargeted aligned BAM files. Inconclusive mpileups, due to different indexing or complex variant calls, were checked manually, and were excluded if read-end or -start marker was present in the mpileup sequence. The resulting mpileup percentage provided the final VAF for our variant calls and was used in all subsequent analyses.

*Quality control multiple-timepoint dataset*

Our multiple-timepoint dataset was subjected to the same quality filtering pipeline, with exception of the run-specific threshold in STEP3 as multiple timepoints of the same individuals constituted one run. The final output from STEP6 was used to trace CHDM calls per individual over all of its available timepoints, to allow most sensitive detection of the same CHDMs appearance at previous timepoints. However, as these supplemental mpileups were not initially identified as CHDMs by JSI Medical Systems Sequence Pilot, our two-fold CHDM detection approach is violated. We therefore added a level of stringency to the parsing of multiple-timepoint mpileups in terms of 1) a position-based coverage threshold of ≥500x, and 2) a minimum alternative allele count threshold of ≥3.

**Statistical Analyses**

All analyses were performed in R version 3.6.1 (R Core Team, URL https://www.R-project.org/), p-values <0.05 were considered statistically significant. Differences in categorical parameters were assessed primarily by means of Chi-square tests, or for expected frequencies <5 by means of Fisher's Exact tests. Differences in continuous parameters were primarily assessed by means of Wilcoxon-rank sum tests.

We created two logistic regression models for our detected single-timepoint mutations using either 1) CHDM prevalence (both small and large clones), or 2) CHIP prevalence (large clones only) as dependent variable. Model fit was assessed by means of a givitiCalibrationBelt-plot using the package givitiR. Parameters of logistic regression models can be found in *File S3, tables 5a and 5b*. The effect of age on clone size was assessed by means of a linear regression model for log-transformed VAF, the resulting parameters can be found in *File S3, table 5c***.**

To determine the effect of age on clone growth in our multiple-timepoint dataset, we first selected the most important trajectory per individual. We then created a mixed linear model (MLM) with random intercept and slope using the nlme package. This model was compared an MLM with random intercept and fixed slope, and an MLM with both fixed parameters by means of the anova() function from the stats package. Details on these model comparisons can be found in *File S3, table 7b*. Finally, to determine whether clinical correlations could underlie differences in speed of clonal outgrowth, we correlated the individual effect estimates from our MLM to aggregated clinical parameters. Specifically, for each parameter the average of baseline, 2- and 10 year follow-up was computed, after which we computed the Spearman correlation coefficient by means of the rcorr() function from the package Hmisc.

All figures were generated in R using a variety of packages (dplyr, reshape2, ggplot2, ggpubr, tidyverse, ggpmisc, rcompanion, ggbeeswarm, ggrepel, ggridges, ggiraphExtra, plotrix, ggallin), after which they were optimized in Adobe Illustrator version 23.11.1 (Adobe Inc.)

***Results:***

**CHDM characteristics and comparison to well-established driver mutations**

As expected and in line with previous reports, *DNMT3A* was the most frequently mutated gene in both our single- and multiple-timepoint dataset (*File S5*). We also identified several loss of function (LoF) mutations in *TET2* and *ASXL1*, and frequently recurring (*i.e.* identified each at least five times in the single-timepoint data of our cohort) missense mutations in *GNAS* [p.(Cys843Arg) and p.(Arg844His)], *GNB1* [p.(Lys57Glu)], *TP53* [p.(Arg174Gly)], and *JAK2* [p.(Val617Phe)]. The majority of CHDMs in the single-timepoint dataset involved the same genes as the growing clones from the multiple-timepoint dataset (*File S5, figs a and b*). The amino acid residues most frequently affected by CHDMs in the single-timepoint dataset were similarly enriched in trajectories that grew in our multiple-timepoint dataset (*File S5, fig. c*).

Our full gene coverage of *DNMT3A* allowed us to examine the location of CHDMs in this gene. We observed that *DNMT3A* missense mutations detected in our single-timepoint dataset clustered around the three known protein domains: PWWP, ADD and SAM-dependent MTase C5-type (*File S5, fig. d*). In our 20-year longitudinal dataset, all LoF mutations in *DNMT3A* were defined as traceable trajectories apart from one late-appearing clone, whereas one third of *DNMT3A* missense mutations were classified as events and two thirds as trajectories (*File S5, fig. e*). Of all the LoF mutations in *DNMT3A*, five out of six early stops or frameshifts (before amino acid 432) were classified as growing (four trajectories and one late-appearing clone), while none of the six later LoF mutations fit this classification. Figure f in *File S5* shows that the clustering of CHDMs in *DNMT3A* previously reported in the literature is comparable to that in both of our datasets, substantiating their role as established driver mutations.

The majority of mutations identified in this study have previously been described (*File S3, table 3a*). Of all the 273 detected single-timepoint mutations, 226 (82.8%) were identical to previously identified CHDMs from the literature or showed an LoF mutation in *DNTM3A, TET2* or *ASXL1*. Of those 226, 61.9% were well-established drivers (≥5 counts in literature) and 10.3% were new substitutions at previously described amino acid positions;

the remaining 7.0%, corresponding to 5 different mutations, were therefore novel candidate CHDMs (*File S3, table 3b*).

Of all the 115 mutations identified in the multiple-timepoint measurements, 71 (61.7%) have been reported in the literature or represent LoF mutations in *DNMT3A, TET2* or *ASXL1,* this refers to 39 (73.58%) when only counting different mutations. Amongst the CHDMs classified as events (*i.e.* mutations only seen at one or two of the timepoints), the overlap with the literature was lowest (12/38; 31.6%). In contrast, the overlap was higher for all CHDMs classified as trajectories: static trajectories (27/42; 64.3%) shrinking trajectories (5/5; 100%) and growing trajectories (27/30; 90.0%) (*File S3, table 3c*).

**References**

1. Sjostrom L, Narbro K, Sjostrom CD, Karason K, Larsson B, Wedel H, Lystig T, Sullivan M, Bouchard C, Carlsson B, Bengtsson C, Dahlgren S, Gummesson A, Jacobson P, Karlsson J, Lindroos AK, Lonroth H, Naslund I, Olbers T, Stenlof K, Torgerson J, Agren G, Carlsson LM, Swedish Obese Subjects S. Effects of bariatric surgery on mortality in Swedish obese subjects. N Engl J Med 2007;**357**(8):741-52.
2. Acuna-Hidalgo R, Sengul H, Steehouwer M, van de Vorst M, Vermeulen SH, Kiemeney L, Veltman JA, Gilissen C, Hoischen A. Ultra-sensitive Sequencing Identifies High Prevalence of Clonal Hematopoiesis-Associated Mutations throughout Adult Life. Am J Hum Genet 2017;**101**(1):50-64.
3. Boyle EA, O'Roak BJ, Martin BK, Kumar A, Shendure J. MIPgen: optimized modeling and design of molecular inversion probes for targeted resequencing. Bioinformatics 2014;**30**(18):2670-2.
4. Li H, Durbin R. Fast and accurate short read alignment with Burrows-Wheeler transform. Bioinformatics 2009;**25**(14):1754-60.
5. Eijkelenboom A, Kamping EJ, Kastner-van Raaij AW, Hendriks-Cornelissen SJ, Neveling K, Kuiper RP, Hoischen A, Nelen MR, Ligtenberg MJ, Tops BB. Reliable Next-Generation Sequencing of Formalin-Fixed, Paraffin-Embedded Tissue Using Single Molecule Tags. J Mol Diagn 2016;**18**(6):851-863.
6. Weren RD, Mensenkamp AR, Simons M, Eijkelenboom A, Sie AS, Ouchene H, van Asseldonk M, Gomez-Garcia EB, Blok MJ, de Hullu JA, Nelen MR, Hoischen A, Bulten J, Tops BB, Hoogerbrugge N, Ligtenberg MJ. Novel BRCA1 and BRCA2 Tumor Test as Basis for Treatment Decisions and Referral for Genetic Counselling of Patients with Ovarian Carcinomas. Hum Mutat 2017;**38**(2):226-235.
7. Li H, Handsaker B, Wysoker A, Fennell T, Ruan J, Homer N, Marth G, Abecasis G, Durbin R, Genome Project Data Processing S. The Sequence Alignment/Map format and SAMtools. Bioinformatics 2009;**25**(16):2078-9.
