## Supplementary FileS2 (figure 1 in supplement) for "Expansion of mutation-driven haematopoietic clones is associated with insulin resistance and low HDL-cholesterol in individuals with obesity"

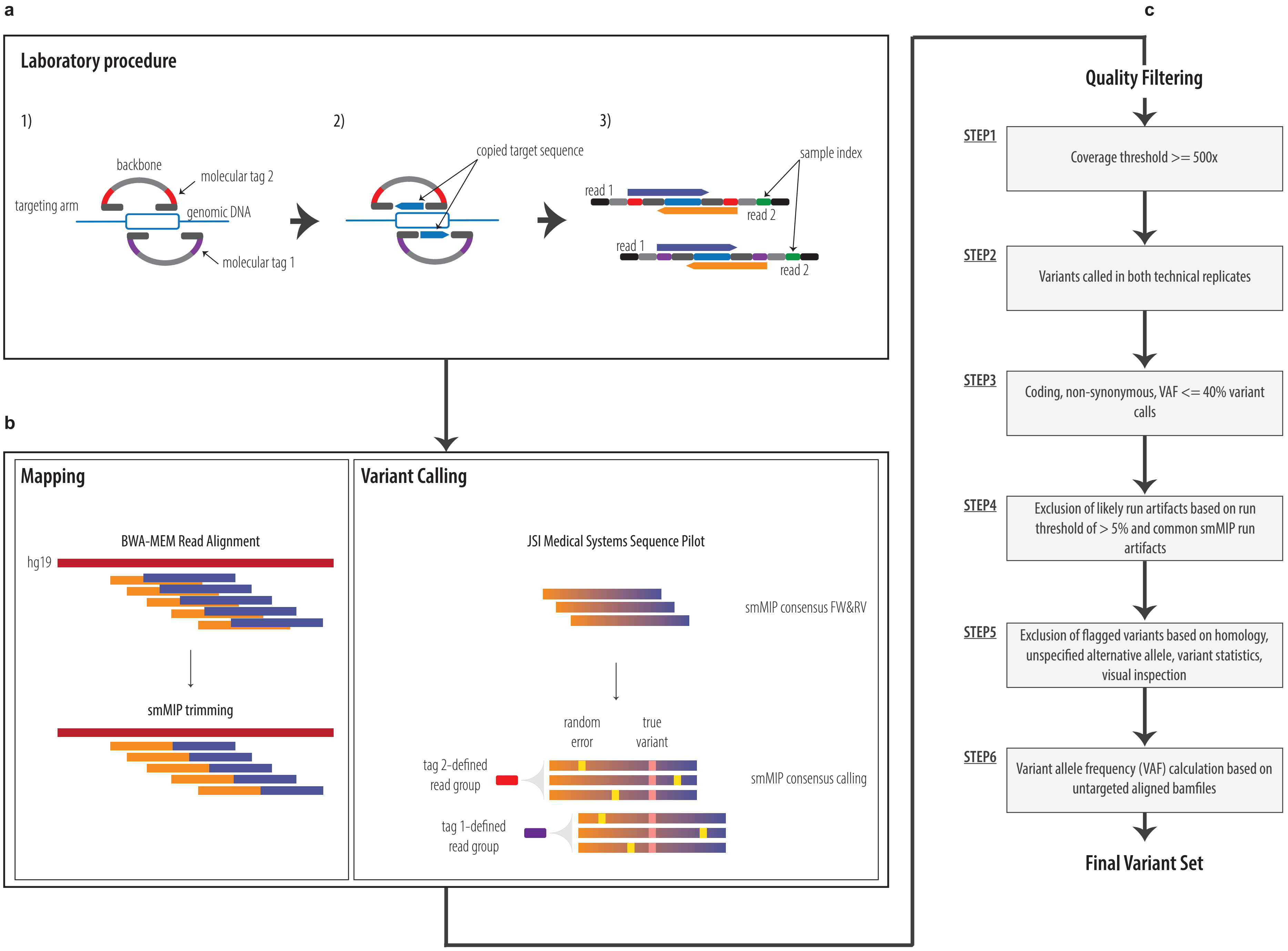
a
Laboratory procedure
1)
backbone
molecular tag 2
targeting arm
genomic DNA
molecular tag 1
2)
copied target sequence
3)
sample index
read 1
read 2
read 1
read 2
b
Mapping
BWA-MEM Read Alignment
hg19
smMIP trimming
Variant Calling
JSI Medical Systems Sequence Pilot
smMIP consensus FW&RV
random error
true variant
tag 2-defined read group
smMIP consensus calling
tag 1-defined read group
c
Quality Filtering
STEP1
Coverage threshold >= 500x
STEP2
Variants called in both technical replicates
STEP3
Coding, non-synonymous, VAF <= 40% variant calls
STEP4
Exclusion of likely run artifacts based on run threshold of > 5% and common smMIP run artifacts
STEP5
Exclusion of flagged variants based on homology, unspecified alternative allele, variant statistics, visual inspection
STEP6
Variant allele frequency (VAF) calculation based on untargeted aligned bamfiles
Final Variant Set

***File S2.*** **smMIP workflow to further boost sensitivity for low-level somatic mutation detection. (a** - adapted from Hiatt et. al. 2013**)** smMIP insert-sizes were shortened to 54nt to enable full forward and reverse read coverage (double sequencing of each insert), target-sequences were generally targeted by at least two independent smMIP-probes (double tiling), and each smMIP capture underwent two idependently barcoded PCR-reactions (double PCR replicates). Sequecing was performed using the Illumina NextSeq 500 system. **(b** - adapted from Hiatt et. al. 2013**)** Raw sequencing data was converted to FastQ-files after which two independent data processing strategies were applied. Mapping: FastQ-file reads were aligned to the reference genome (Hg19) with BWA-MEM after which the overlap between Forward (FW) and Reverse (RV) reads was trimmed off (smMIP trimming). Variant Calling: FastQ-files were imported in JSI Medical Systems Sequence Pilot in which first a consensus between FW and RV reads is determined (smMIP consensus FW&RV), after which tag-defined read groups enable smMIP consensus calling. **(c)** The aligned reads and resulting variant calls are then subjected to a stringent quality filtering pipeline. First, individuals with an average sequencing depth <500x are excluded (STEP1). Second, only variants called in both technical replicates are kept (STEP2). Third, coding, non-synonymous variant calls with a Variant Allele Frequency (VAF) <= 40% are kept (STEP3). Fourth, likely run-specific artifacts (variant calls in > 5% of samples per run) are excluded (STEP4). Fifth, exclusion of variants based on homology, unspecified alternative allele, variant statistics, and visual inspection (STEP5). And finally, for each variant position we generated mpileups based on the aligned reads to determine the final VAF for each CHDM (STEP6).
