## Supplementary material for "Expansion of mutation-driven haematopoietic clones is associated with insulin resistance and low HDL-cholesterol in individuals with obesity": Supplentary FileS4 (figure 2 in supplement)

a

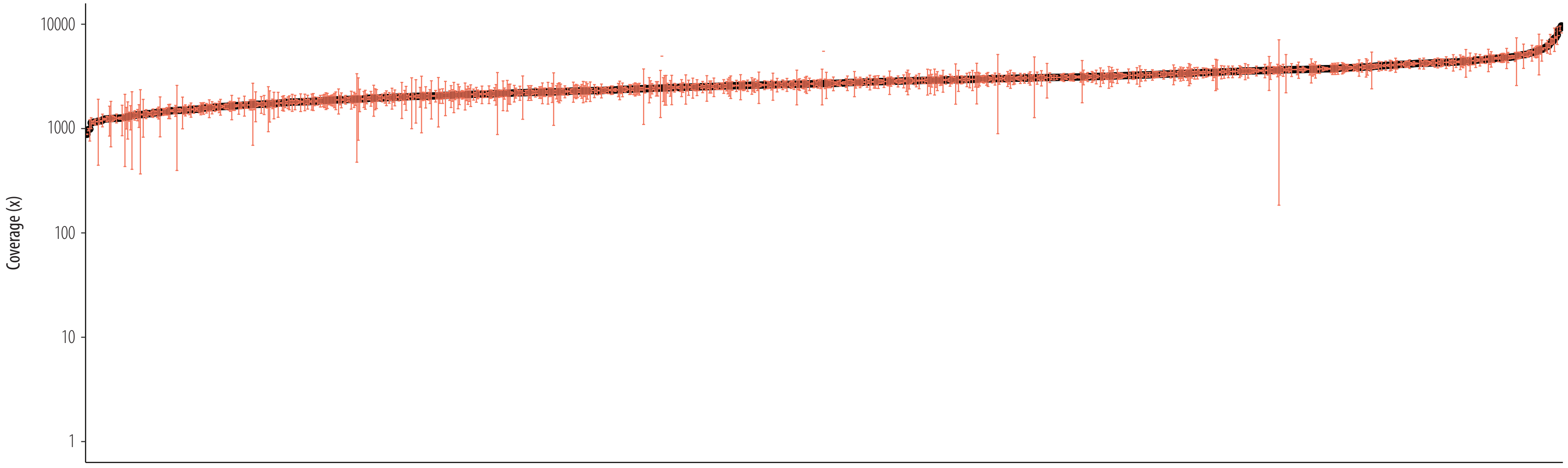


b

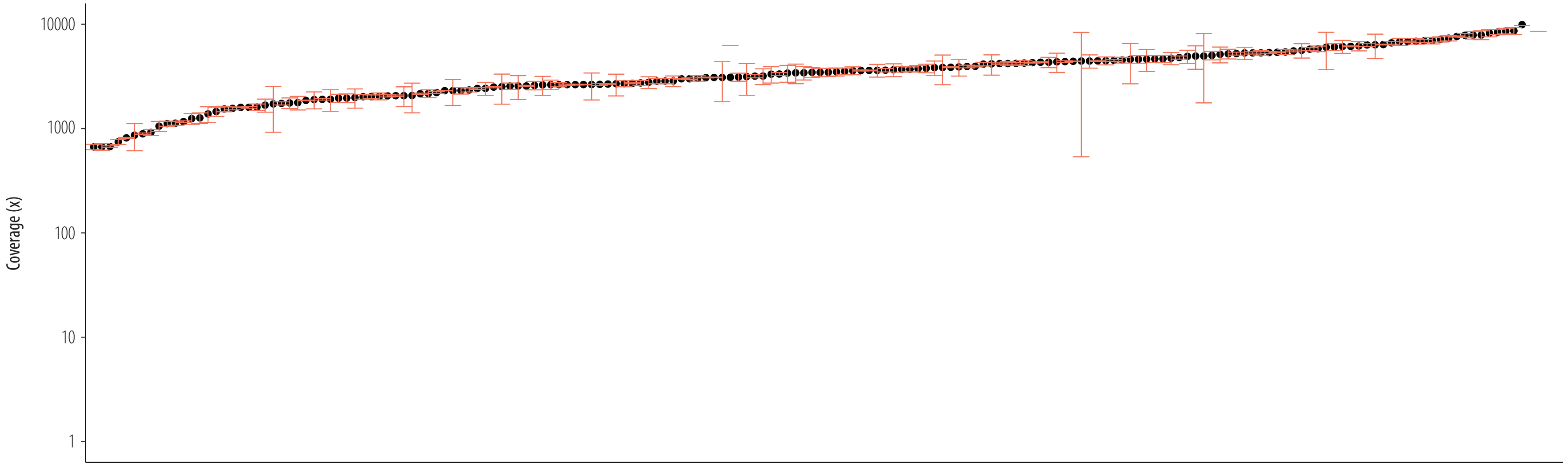


***File S4.*** **smMIP coverage of single- and multiple-timepoint datasets. (a)** Average smMIP coverage over the entire CH-panel for both technical replicates per individual in the single-timepoint dataset (N=1050). **(b)** Average smMIP coverage over the entire CH-panel for both technical replicates per DNA sample in the multiple-timepoint dataset (N DNA samples=180). In both **(a)** and **(b)** the error bars represent the standard deviation.
