## Supplementary FileS5 (figure 3 in supplement) for "Expansion of mutation-driven haematopoietic clones is associated with insulin resistance and low HDL-cholesterol in individuals with obesity"

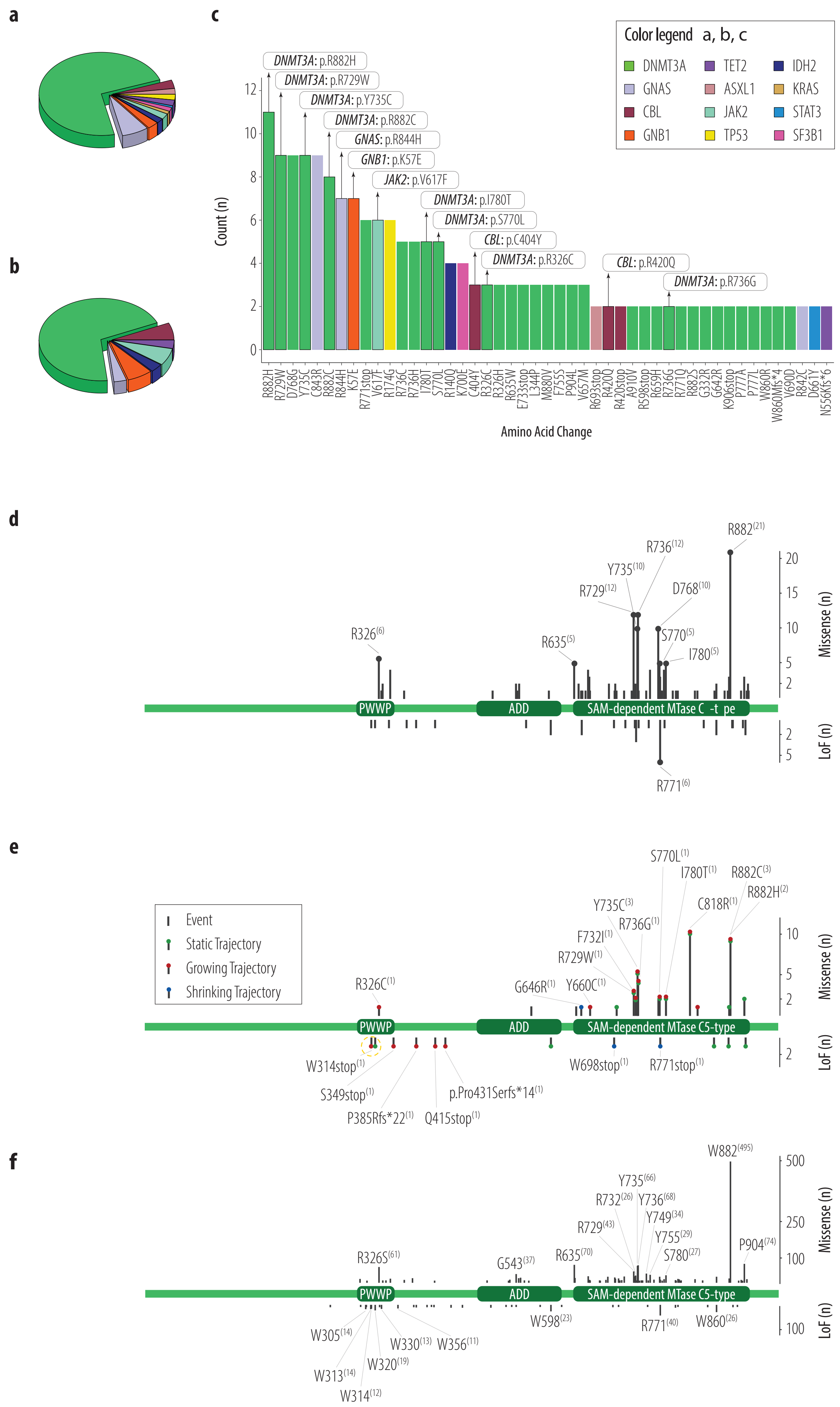

a
b
c
Color legend a, b, c
DNMT3A
GNAS
CBL
GNB1
TET2
ASXL1
JAK2
TP53
IDH2
KRAS
STAT3
SF3B1
DNMT3A: p.R882H
DNMT3A: p.R729W
DNMT3A: p.Y735C
DNMT3A: p.R882C
GNAS: p.R844H
GNB1: p.K57E
JAK2: p.V617F
DNMT3A: p.I780T
DNMT3A: p.S770L
CBL: p.C404Y
DNMT3A: p.R326C
CBL: p.R420Q
DNMT3A: p.R736G
Count (n)
0
2
4
6
8
10
12
R882H R729W D768G Y735C C843R R882C R844H K57E R771stop V617F R174G R736C R736H I780T S770L R140Q K700E C404Y R326C R326H R635W E733stop L344P M880V F755S P904L V657M R693stop R420Q R420stop A910V R598stop R659H R736G R771Q R882S G332R G642R K906stop P777A P777L W860R W860Mfs*4 V690D R842C D661Y N556Kfs*6
Amino Acid Change
d
R882(21)
R736(12)
Y735(10)
R729(12)
D768(10)
R326(6)
R635(5)
S770(5)
I780(5)
PWWP
ADD
SAM-dependent MTase C -t pe
R771(6)
Missense (n)
LoF (n)
20
15
10
5
2
e
Event
Static Trajectory
Growing Trajectory
Shrinking Trajectory
S770L(1)
I780T(1)
R882C(3)
R882H(2)
Y735C(3)
C818R(1)
R736G(1)
F732I(1)
R729W(1)
R326C(1)
G646R(1)
Y660C(1)
PWWP
ADD
SAM-dependent MTase C5-type
W314stop(1)
S349stop(1)
P385Rfs*22(1)
Q415stop(1)
p.Pro431Serfs*14(1)
W698stop(1)
R771stop(1)
Missense (n)
LoF (n)
10
5
2
f
W882(495)
Y735(66)
R732(26)
Y736(68)
Y749(34)
R729(43)
Y755(29)
S780(27)
P904(74)
R326S(61)
G543(37)
R635(70)
PWWP
ADD
SAM-dependent MTase C5-type
W305(14)
W313(14)
W314(12)
W320(19)
W330(13)
W356(11)
W598(23)
R771(40)
W860(26)
Missense (n)
LoF (n)
500
250
100

***File S5.*** **CHDMs identified in the single- and multiple timepoint datasets and comparison to literature. (a)** The gene distribution of identified CHDMs in the single-timepoint dataset (total number of CHDMs=273). **(b)** The gene distribution of growing CHDM trajectories in the multiple-timepoint dataset (total number of growing trajectories=30). **(c)** The number of CHDMs resulting in specific amino acid changes in the single-timepoint dataset at the protein level. Labels indicate CHDMs that have been defined as growing trajectories in our multiple-timepoint dataset. **(d)** Amino acid changes caused by DNMT3A mutations in our single-timepoint dataset annotated on a schematic visualization of the DNMT3A protein. Mutations affecting the same amino acid residue five or more times have been annotated and the label specifies the reference amino acid, position and number of mutations in superscript. Labels: R326$^{(6)}$ = R326H (n=3), R326C (n=3); R635$^{(5)}$ = R635W (n=3), R635Q (n=1), R635L (n=1); R729$^{(12)}$ = R729W (n=9), R729Q (n=1), R729G (n=1), R729L (n=1); Y735$^{(10)}$ = Y735C (n=9), Y735S (n=1); R736$^{(12)}$ = R736C (n=5), R736H (n=5), R736G (n=2); D768$^{(10)}$ = D768G (n=9), D768V (n=1); R771$^{(6)}$ = R771stop (n=6); R882$^{(21)}$ = R882H (n=11), R882C (n=8), R882S (n=2). **(e)** Amino acid changes caused by DNMT3A mutations in our multiple-timepoint dataset annotated on a schematic visualization of DNMT3A protein. CHDMs that have been categorized as trajectories have been given a coloured dot: red for growing (with a dashed yellow circle for 'late-appearing clone'), green for static and blue for shrinking. Labels constitute the reference amino acid in single letter code, position, and alternative amino acid, and are provided only for growing and shrinking trajectories (as represented by the number in superscript), and not for static trajectories or events (with the exception of 'late-appearing clone'). **(f)** DNMT3A mutations (amino acid changes) observed at least five times in literature annotated on a schematic visualization of DNMT3A protein. Missense mutations observed >25 times, and LoF mutations observed >10 times have been annotated and the label specifies the reference amino acid, position and number of calls in superscript. For a complete table of all CHDMs observed in literature, see File S3, table 3a. DNMT3A protein annotation source: https://www.uniprot.org/uniprot/Q9Y6K1.
