## Supplementary FileS6 (figure 4 in supplement) for "Expansion of mutation-driven haematopoietic clones is associated with insulin resistance and low HDL-cholesterol in individuals with obesity"

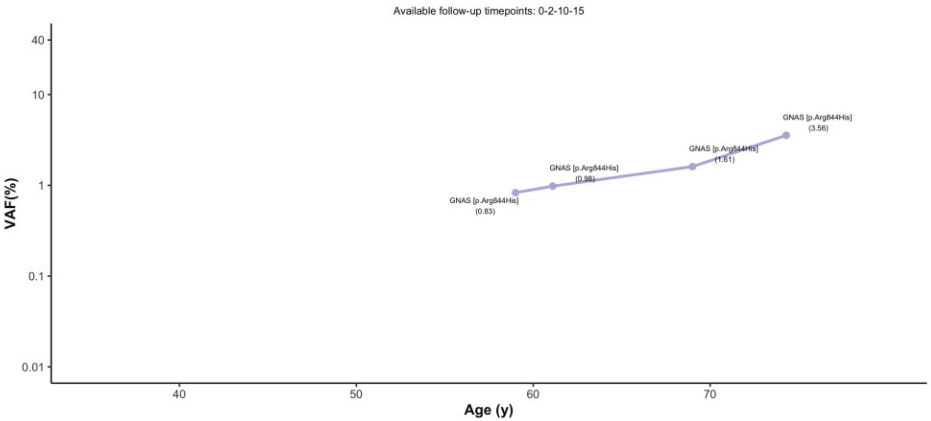
Available follow-up timepoints: 0-2-10-15
40
10
1
0.1
0.01
VAF(%)
40
50
60
70
Age (y)
GNAS [p.Arg844His]
(0.83)
GNAS [p.Arg844His]
(0.98)
GNAS [p.Arg844His]
(1.61)
GNAS [p.Arg844His]
(3.56)

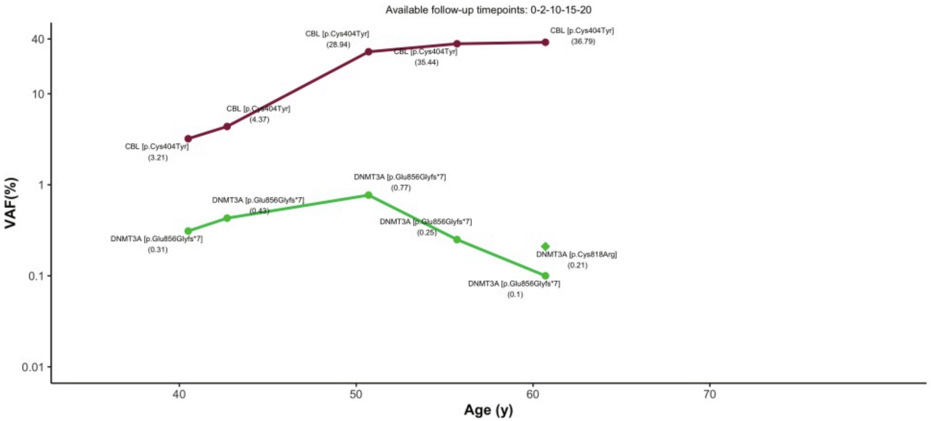
Available follow-up timepoints: 0-2-10-15-20
40
10
1
0.1
0.01
VAF(%)
40
50
60
70
Age (y)
CBL [p.Cys404Tyr]
(3.21)
CBL [p.Cys404Tyr]
(4.37)
CBL [p.Cys404Tyr]
(28.94)
CBL [p.Cys404Tyr]
(35.44)
CBL [p.Cys404Tyr]
(36.79)
DNMT3A [p.Glu856Glyfs*7]
(0.31)
DNMT3A [p.Glu856Glyfs*7]
(0.43)
DNMT3A [p.Glu856Glyfs*7]
(0.77)
DNMT3A [p.Glu856Glyfs*7]
(0.25)
DNMT3A [p.Glu856Glyfs*7]
(0.1)
DNMT3A [p.Cys818Arg]
(0.21)

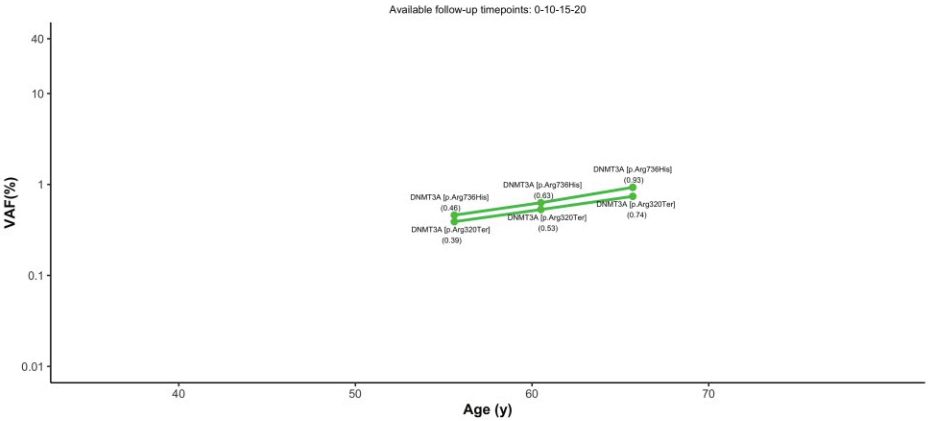
Available follow-up timepoints: 0-10-15-20
40
10
1
0.1
0.01
VAF(%)
40
50
60
70
Age (y)
DNMT3A [p.Arg736His]
(0.46)
DNMT3A [p.Arg736His]
(0.63)
DNMT3A [p.Arg736His]
(0.93)
DNMT3A [p.Arg320Ter]
(0.39)
DNMT3A [p.Arg320Ter]
(0.53)
DNMT3A [p.Arg320Ter]
(0.74)

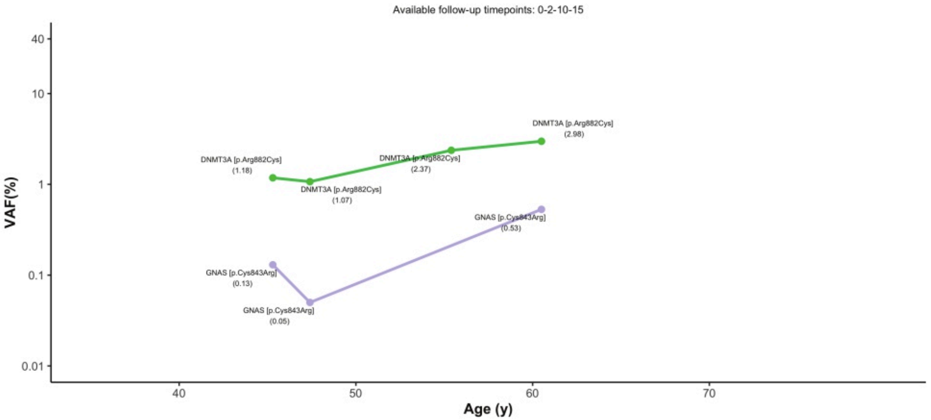
Available follow-up timepoints: 0-2-10-15
40
10
1
0.1
0.01
VAF(%)
40
50
60
70
Age (y)
DNMT3A [p.Arg882Cys]
(1.18)
DNMT3A [p.Arg882Cys]
(1.07)
DNMT3A [p.Arg882Cys]
(2.37)
DNMT3A [p.Arg882Cys]
(2.98)
GNAS [p.Cys843Arg]
(0.13)
GNAS [p.Cys843Arg]
(0.05)
GNAS [p.Cys843Arg]
(0.53)

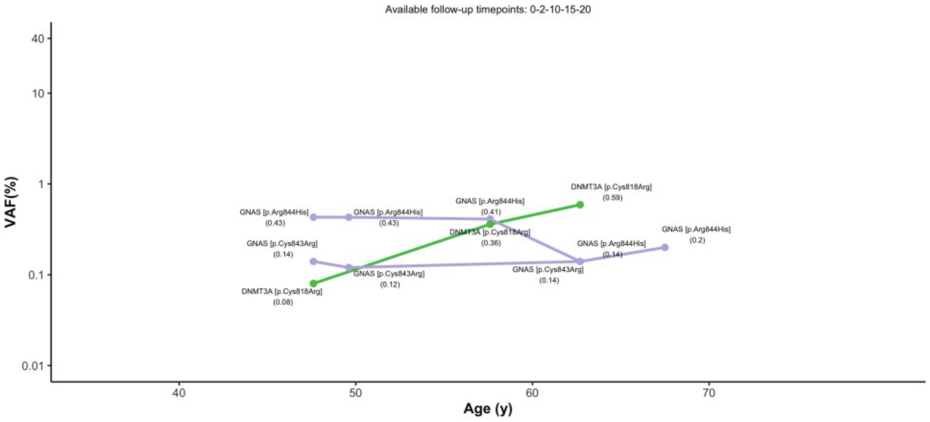
Available follow-up timepoints: 0-2-10-15-20
VAF(%)
Age (y)
GNAS [p.Arg844His] (0.43)
GNAS [p.Arg844His] (0.43)
GNAS [p.Arg844His] (0.41)
DNMT3A [p.Cys818Arg] (0.59)
GNAS [p.Arg844His] (0.2)
DNMT3A [p.Cys818Arg] (0.36)
GNAS [p.Arg844His] (0.14)
GNAS [p.Cys843Arg] (0.14)
GNAS [p.Cys843Arg] (0.12)
GNAS [p.Cys843Arg] (0.14)
DNMT3A [p.Cys818Arg] (0.08)

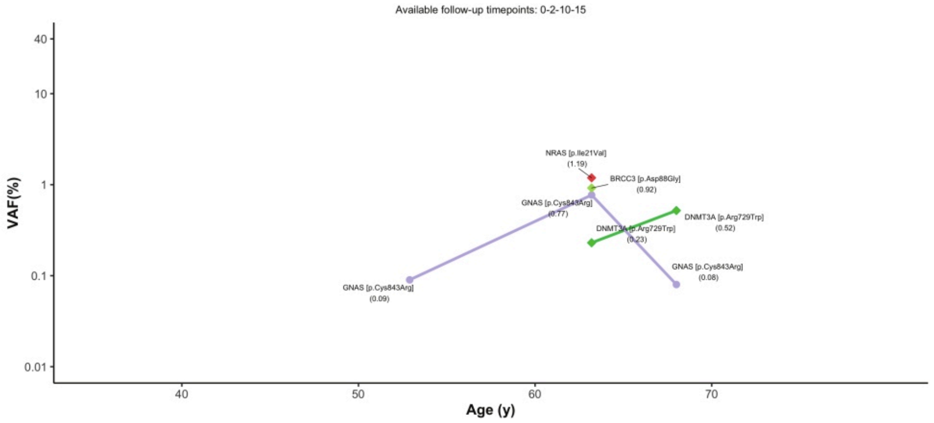
Available follow-up timepoints: 0-2-10-15
VAF(%)
Age (y)
NRAS [p.Ile21Val] (1.19)
BRCC3 [p.Asp88Gly] (0.92)
GNAS [p.Cys843Arg] (0.77)
DNMT3A [p.Arg729Trp] (0.52)
DNMT3A [p.Arg729Trp] (0.23)
GNAS [p.Cys843Arg] (0.08)
GNAS [p.Cys843Arg] (0.09)

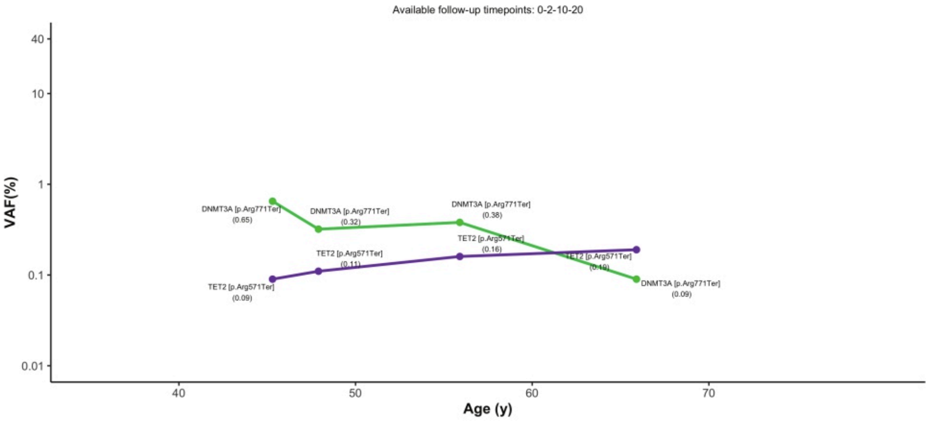
Available follow-up timepoints: 0-2-10-20
VAF(%)
Age (y)
DNMT3A [p.Arg771Ter] (0.65)
DNMT3A [p.Arg771Ter] (0.32)
DNMT3A [p.Arg771Ter] (0.38)
TET2 [p.Arg571Ter] (0.16)
TET2 [p.Arg571Ter] (0.11)
TET2 [p.Arg571Ter] (0.19)
TET2 [p.Arg571Ter] (0.09)
DNMT3A [p.Arg771Ter] (0.09)

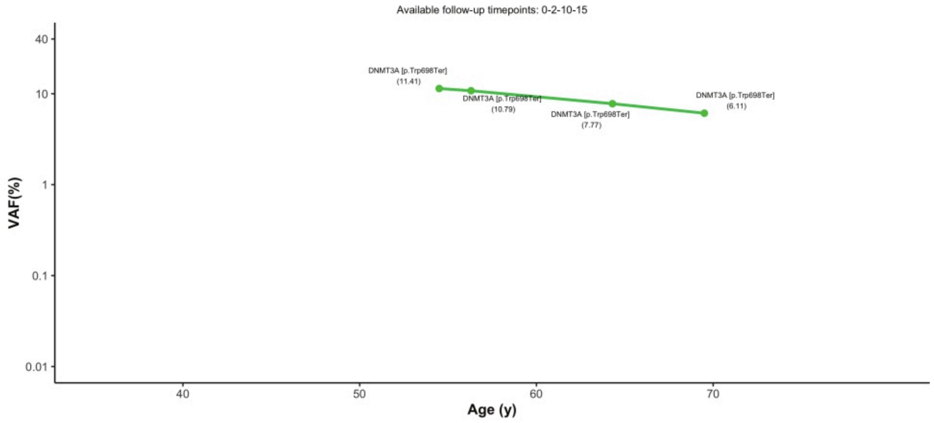
Available follow-up timepoints: 0-2-10-15
VAF(%)
Age (y)
DNMT3A [p.Trp698Ter] (11.41)
DNMT3A [p.Trp698Ter] (10.79)
DNMT3A [p.Trp698Ter] (7.77)
DNMT3A [p.Trp698Ter] (6.11)

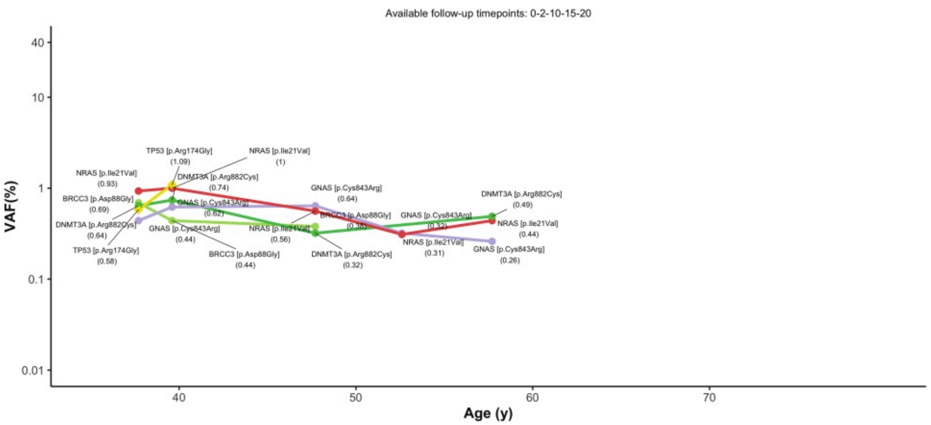

Available follow-up timepoints: 0-2-10-15-20
VAF(%)
Age (y)
40
10
1
0.1
0.01
40
50
60
70
TP53 [p.Arg174Gly] (1.09)
NRAS [p.Ile21Val] (1)
NRAS [p.Ile21Val] (0.93)
DNMT3A [p.Arg882Cys] (0.74)
BRCC3 [p.Asp88Gly] (0.69)
GNAS [p.Cys843Arg] (0.62)
DNMT3A [p.Arg882Cys] (0.64)
GNAS [p.Cys843Arg] (0.44)
TP53 [p.Arg174Gly] (0.58)
BRCC3 [p.Asp88Gly] (0.44)
GNAS [p.Cys843Arg] (0.64)
BRCC3 [p.Asp88Gly] (0.35)
NRAS [p.Ile21Val] (0.56)
DNMT3A [p.Arg882Cys] (0.32)
GNAS [p.Cys843Arg] (0.32)
NRAS [p.Ile21Val] (0.31)
DNMT3A [p.Arg882Cys] (0.49)
NRAS [p.Ile21Val] (0.44)
GNAS [p.Cys843Arg] (0.26)


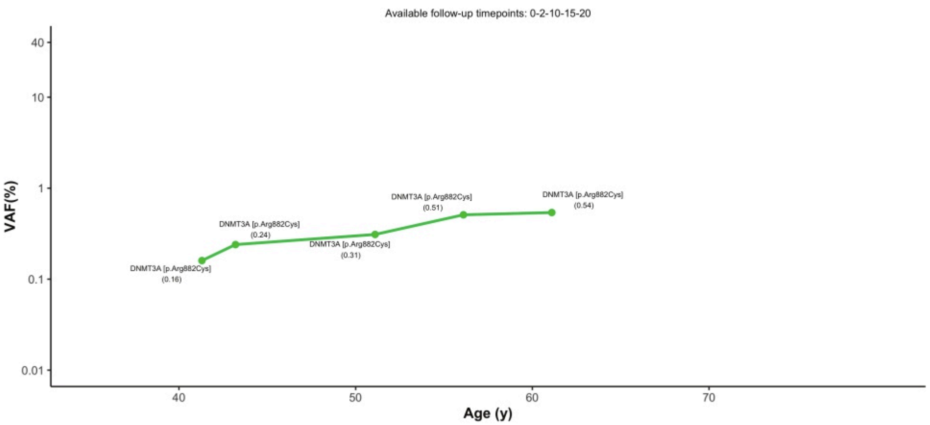

Available follow-up timepoints: 0-2-10-15-20
VAF(%)
Age (y)
40
10
1
0.1
0.01
40
50
60
70
DNMT3A [p.Arg882Cys] (0.16)
DNMT3A [p.Arg882Cys] (0.24)
DNMT3A [p.Arg882Cys] (0.31)
DNMT3A [p.Arg882Cys] (0.51)
DNMT3A [p.Arg882Cys] (0.54)


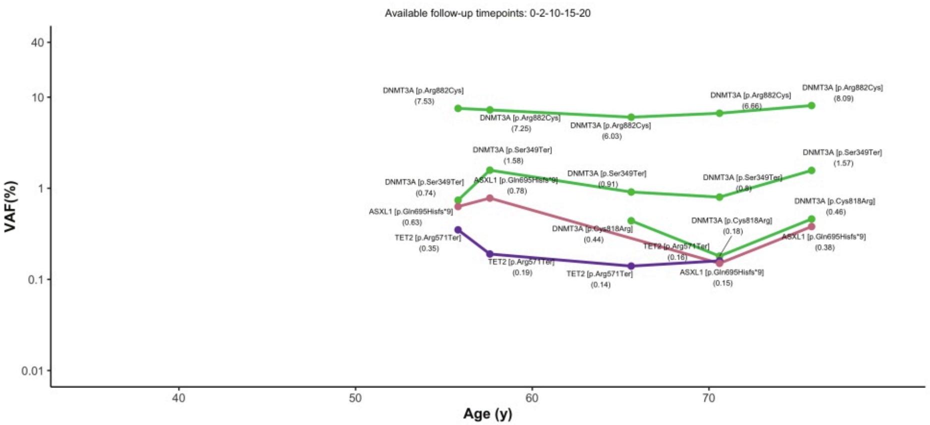

Available follow-up timepoints: 0-2-10-15-20
VAF(%)
Age (y)
40
10
1
0.1
0.01
40
50
60
70
DNMT3A [p.Arg882Cys] (7.53)
DNMT3A [p.Arg882Cys] (7.25)
DNMT3A [p.Arg882Cys] (6.03)
DNMT3A [p.Arg882Cys] (6.66)
DNMT3A [p.Arg882Cys] (8.09)
DNMT3A [p.Ser349Ter] (1.58)
DNMT3A [p.Ser349Ter] (0.74)
ASXL1 [p.Gln695Hisfs*9] (0.78)
DNMT3A [p.Ser349Ter] (0.91)
DNMT3A [p.Ser349Ter] (0.8)
DNMT3A [p.Ser349Ter] (1.57)
ASXL1 [p.Gln695Hisfs*9] (0.63)
DNMT3A [p.Cys818Arg] (0.46)
DNMT3A [p.Cys818Arg] (0.18)
DNMT3A [p.Cys818Arg] (0.44)
TET2 [p.Arg571Ter] (0.35)
TET2 [p.Arg571Ter] (0.16)
ASXL1 [p.Gln695Hisfs*9] (0.38)
TET2 [p.Arg571Ter] (0.19)
TET2 [p.Arg571Ter] (0.14)
ASXL1 [p.Gln695Hisfs*9] (0.15)


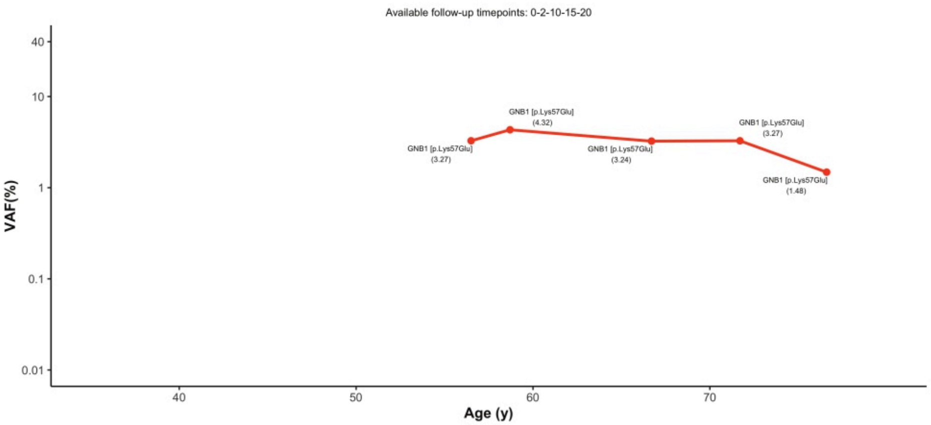

Available follow-up timepoints: 0-2-10-15-20
VAF(%)
Age (y)
40
10
1
0.1
0.01
40
50
60
70
GNB1 [p.Lys57Glu] (4.32)
GNB1 [p.Lys57Glu] (3.27)
GNB1 [p.Lys57Glu] (3.24)
GNB1 [p.Lys57Glu] (3.27)
GNB1 [p.Lys57Glu] (1.48)

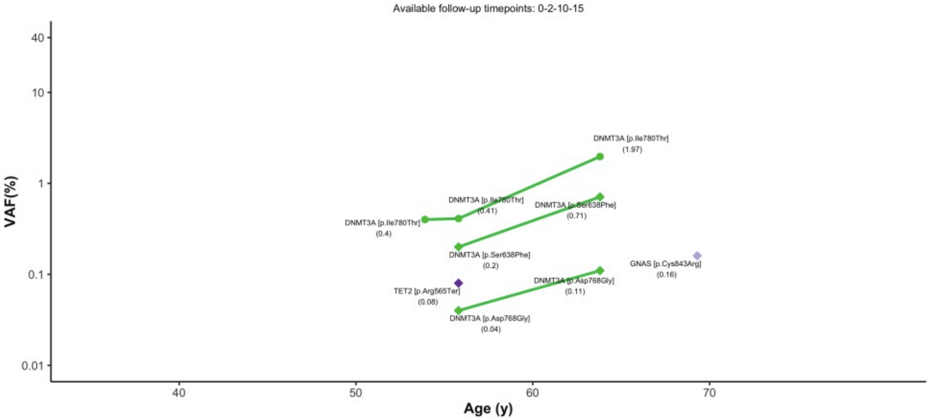
Available follow-up timepoints: 0-2-10-15
DNMT3A [p.Ile780Thr]
(1.97)
DNMT3A [p.Ile780Thr]
(0.41)
DNMT3A [p.Ser638Phe]
(0.71)
DNMT3A [p.Ile780Thr]
(0.4)
DNMT3A [p.Ser638Phe]
(0.2)
GNAS [p.Cys843Arg]
(0.16)
DNMT3A [p.Asp768Gly]
(0.11)
TET2 [p.Arg565Ter]
(0.08)
DNMT3A [p.Asp768Gly]
(0.04)
VAF(%)
Age (y)

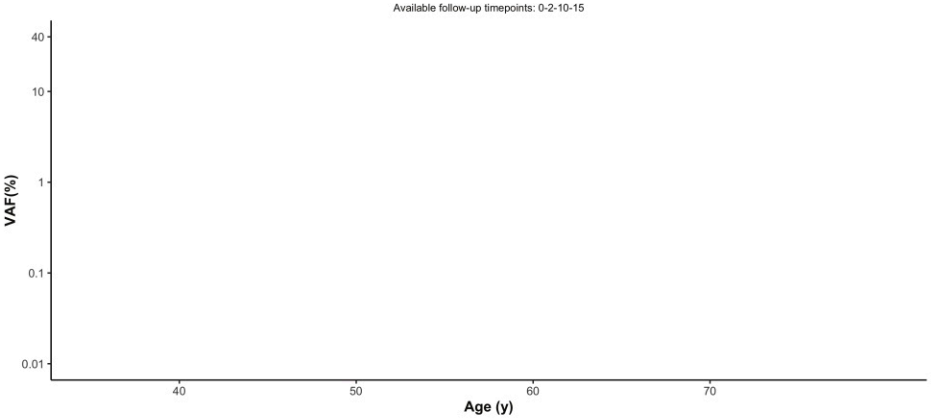
Available follow-up timepoints: 0-2-10-15
VAF(%)
Age (y)

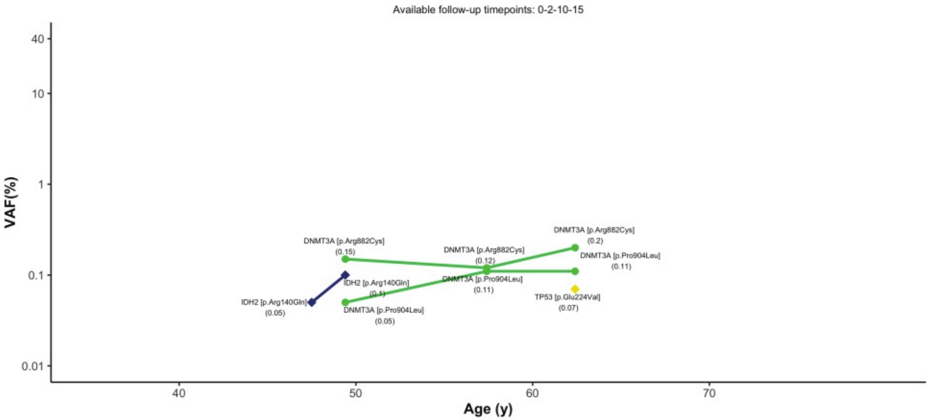
Available follow-up timepoints: 0-2-10-15
DNMT3A [p.Arg882Cys]
(0.2)
DNMT3A [p.Arg882Cys]
(0.15)
DNMT3A [p.Arg882Cys]
(0.12)
DNMT3A [p.Pro904Leu]
(0.11)
IDH2 [p.Arg140Gln]
(0.1)
DNMT3A [p.Pro904Leu]
(0.11)
TP53 [p.Glu224Val]
(0.07)
IDH2 [p.Arg140Gln]
(0.05)
DNMT3A [p.Pro904Leu]
(0.05)
VAF(%)
Age (y)

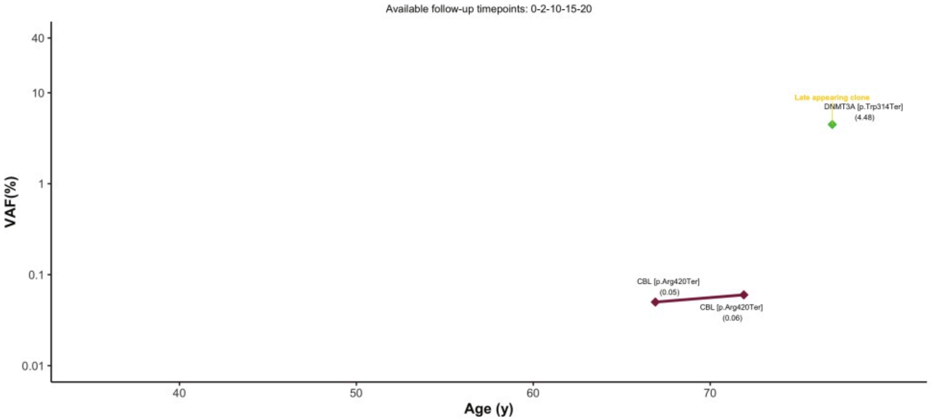
Available follow-up timepoints: 0-2-10-15-20
Late appearing clone
DNMT3A [p.Trp314Ter]
(4.48)
CBL [p.Arg420Ter]
(0.05)
CBL [p.Arg420Ter]
(0.06)
VAF(%)
Age (y)

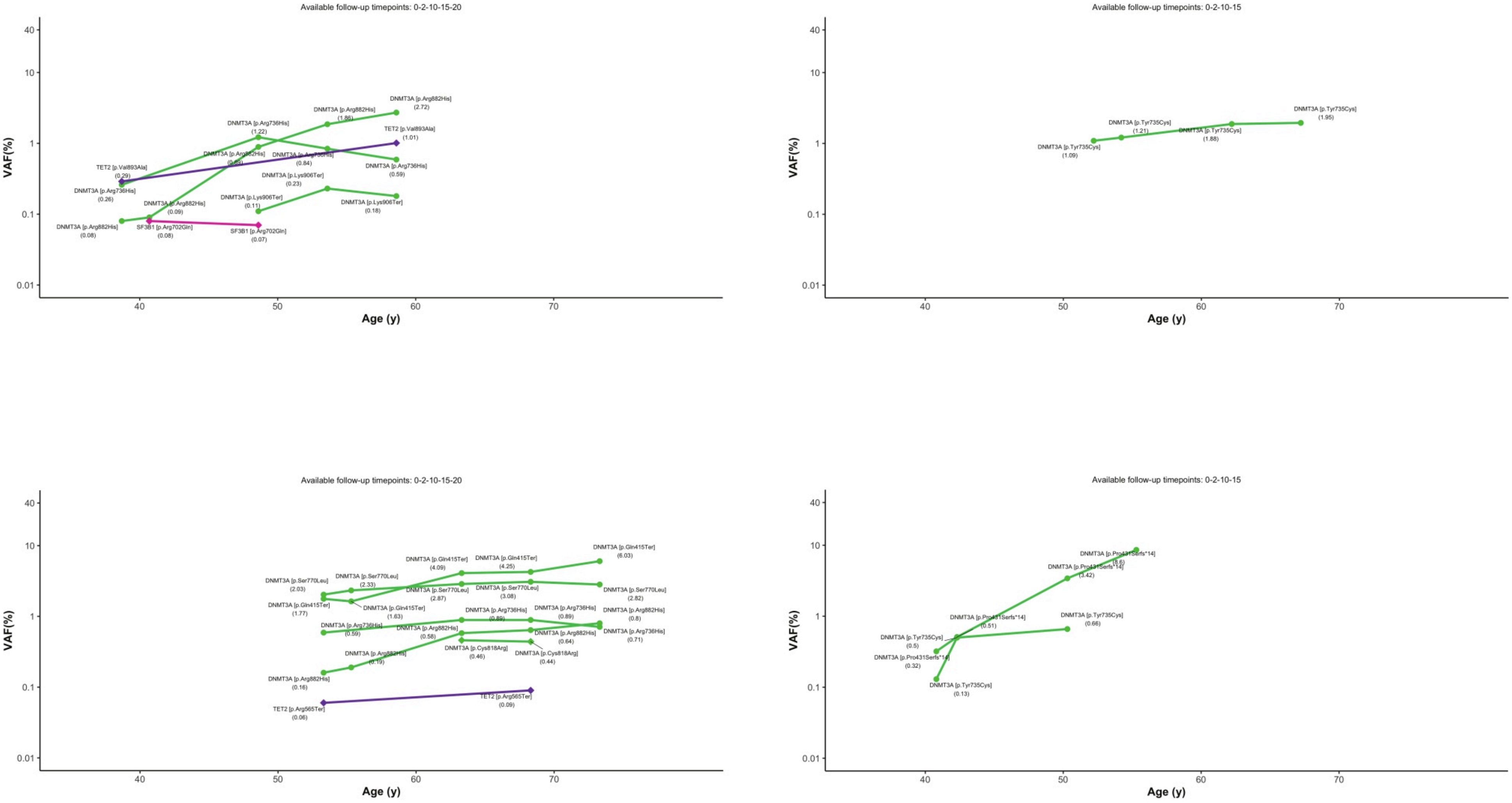
Available follow-up timepoints: 0-2-10-15-20
VAF(%)
Age (y)
DNMT3A [p.Arg882His] (2.72)
DNMT3A [p.Arg882His] (1.86)
DNMT3A [p.Arg736His] (1.22)
TET2 [p.Val893Ala] (1.01)
DNMT3A [p.Arg736His] (0.84)
DNMT3A [p.Arg736His] (0.59)
TET2 [p.Val893Ala] (0.29)
DNMT3A [p.Arg736His] (0.26)
DNMT3A [p.Lys906Ter] (0.23)
DNMT3A [p.Lys906Ter] (0.11)
DNMT3A [p.Lys906Ter] (0.18)
DNMT3A [p.Arg882His] (0.09)
DNMT3A [p.Arg882His] (0.08)
SF3B1 [p.Arg702Gln] (0.08)
SF3B1 [p.Arg702Gln] (0.07)
Available follow-up timepoints: 0-2-10-15
DNMT3A [p.Tyr735Cys] (1.95)
DNMT3A [p.Tyr735Cys] (1.21)
DNMT3A [p.Tyr735Cys] (1.88)
DNMT3A [p.Tyr735Cys] (1.09)
Available follow-up timepoints: 0-2-10-15-20
DNMT3A [p.Gln415Ter] (6.03)
DNMT3A [p.Gln415Ter] (4.09)
DNMT3A [p.Gln415Ter] (4.25)
DNMT3A [p.Ser770Leu] (2.03)
DNMT3A [p.Ser770Leu] (2.33)
DNMT3A [p.Ser770Leu] (2.87)
DNMT3A [p.Ser770Leu] (3.08)
DNMT3A [p.Ser770Leu] (2.82)
DNMT3A [p.Gln415Ter] (1.77)
DNMT3A [p.Gln415Ter] (1.63)
DNMT3A [p.Arg736His] (0.89)
DNMT3A [p.Arg736His] (0.89)
DNMT3A [p.Arg882His] (0.8)
DNMT3A [p.Arg736His] (0.71)
DNMT3A [p.Arg736His] (0.59)
DNMT3A [p.Arg882His] (0.58)
DNMT3A [p.Arg882His] (0.64)
DNMT3A [p.Cys818Arg] (0.46)
DNMT3A [p.Cys818Arg] (0.44)
DNMT3A [p.Arg882His] (0.19)
DNMT3A [p.Arg882His] (0.16)
TET2 [p.Arg565Ter] (0.06)
TET2 [p.Arg565Ter] (0.09)
Available follow-up timepoints: 0-2-10-15
DNMT3A [p.Pro431Serfs*14] (8.6)
DNMT3A [p.Pro431Serfs*14] (3.42)
DNMT3A [p.Pro431Serfs*14] (0.51)
DNMT3A [p.Tyr735Cys] (0.66)
DNMT3A [p.Tyr735Cys] (0.5)
DNMT3A [p.Pro431Serfs*14] (0.32)
DNMT3A [p.Tyr735Cys] (0.13)

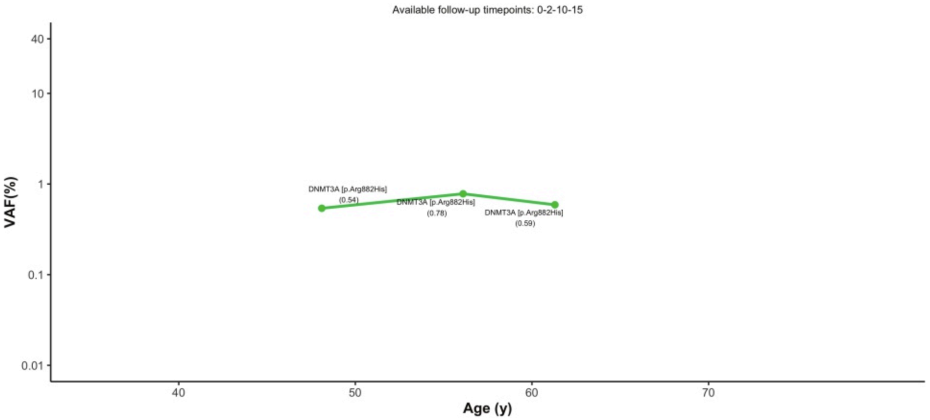
Available follow-up timepoints: 0-2-10-15
40
10
1
0.1
0.01
VAF(%)
40
50
60
70
Age (y)
DNMT3A [p.Arg882His]
(0.54)
DNMT3A [p.Arg882His]
(0.78)
DNMT3A [p.Arg882His]
(0.59)

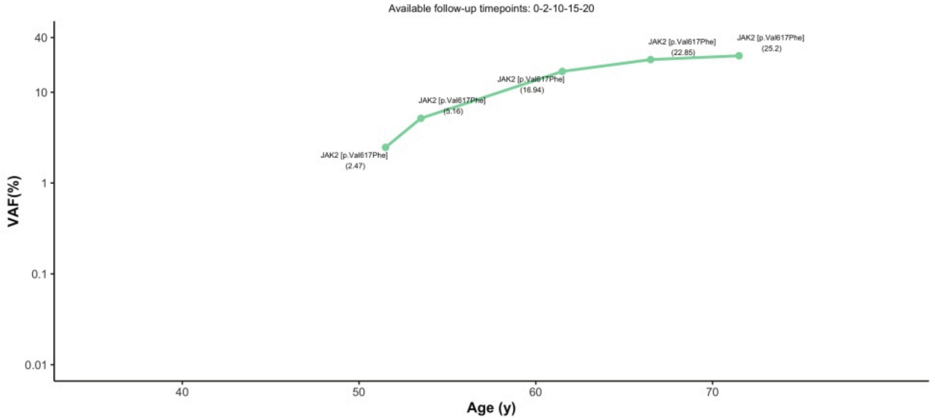
Available follow-up timepoints: 0-2-10-15-20
40
10
1
0.1
0.01
VAF(%)
40
50
60
70
Age (y)
JAK2 [p.Val617Phe]
(2.47)
JAK2 [p.Val617Phe]
(5.16)
JAK2 [p.Val617Phe]
(16.94)
JAK2 [p.Val617Phe]
(22.85)
JAK2 [p.Val617Phe]
(25.2)

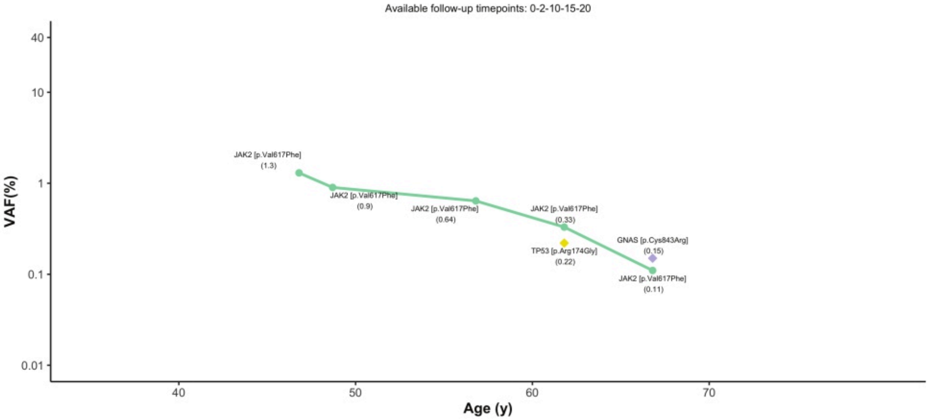
Available follow-up timepoints: 0-2-10-15-20
40
10
1
0.1
0.01
VAF(%)
40
50
60
70
Age (y)
JAK2 [p.Val617Phe]
(1.3)
JAK2 [p.Val617Phe]
(0.9)
JAK2 [p.Val617Phe]
(0.64)
JAK2 [p.Val617Phe]
(0.33)
TP53 [p.Arg174Gly]
(0.22)
GNAS [p.Cys843Arg]
(0.15)
JAK2 [p.Val617Phe]
(0.11)

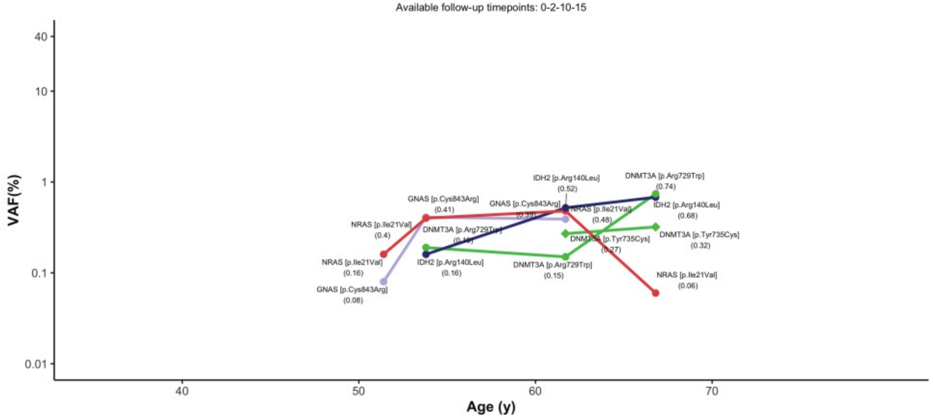
Available follow-up timepoints: 0-2-10-15
40
10
1
0.1
0.01
VAF(%)
40
50
60
70
Age (y)
NRAS [p.Ile21Val]
(0.4)
NRAS [p.Ile21Val]
(0.16)
GNAS [p.Cys843Arg]
(0.08)
GNAS [p.Cys843Arg]
(0.41)
DNMT3A [p.Arg729
IDH2 [p.Arg140Leu]
(0.16)
GNAS [p.Cys843Arg]
IDH2 [p.Arg140Leu]
(0.52)
NRAS [p.Ile21Val]
(0.48)
DNMT3A [p.Arg729Trp]
(0.15)
DNMT3A [p.Tyr735Cys]
(0.27)
DNMT3A [p.Arg729Trp]
(0.74)
IDH2 [p.Arg140Leu]
(0.68)
DNMT3A [p.Tyr735Cys]
(0.32)
NRAS [p.Ile21Val]
(0.06)

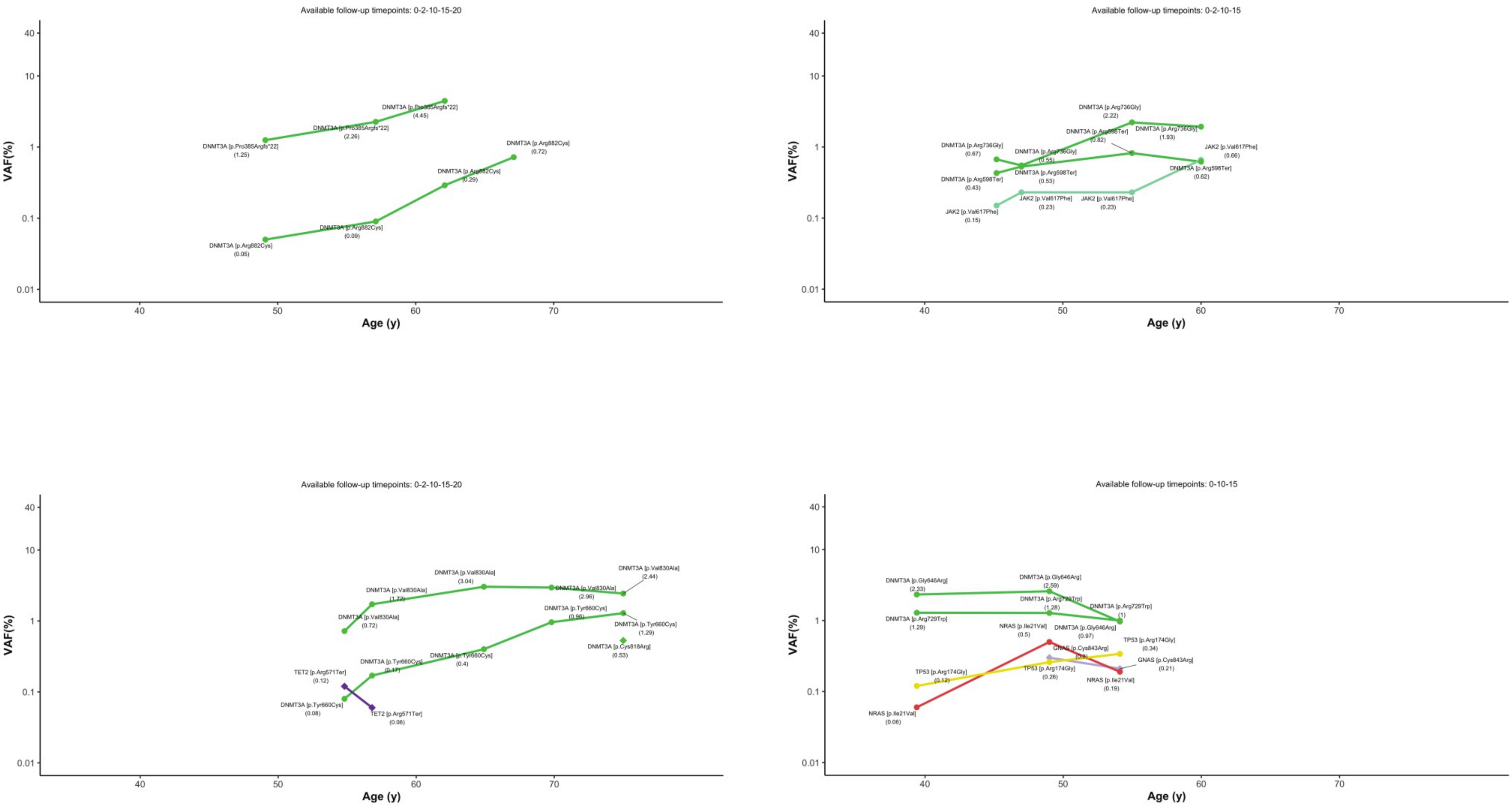
Available follow-up timepoints: 0-2-10-15-20
VAF(%)
Age (y)
DNMT3A [p.Pro385Argfs*22] (1.25)
DNMT3A [p.Pro385Argfs*22] (2.26)
DNMT3A [p.Pro385Argfs*22] (4.45)
DNMT3A [p.Arg882Cys] (0.05)
DNMT3A [p.Arg882Cys] (0.09)
DNMT3A [p.Arg882Cys] (0.29)
DNMT3A [p.Arg882Cys] (0.72)
Available follow-up timepoints: 0-2-10-15
DNMT3A [p.Arg736Gly] (0.67)
DNMT3A [p.Arg736Gly] (0.55)
DNMT3A [p.Arg736Gly] (2.22)
DNMT3A [p.Arg736Gly] (1.93)
DNMT3A [p.Arg598Ter] (0.43)
DNMT3A [p.Arg598Ter] (0.53)
DNMT3A [p.Arg598Ter] (0.82)
DNMT3A [p.Arg598Ter] (0.62)
JAK2 [p.Val617Phe] (0.15)
JAK2 [p.Val617Phe] (0.23)
JAK2 [p.Val617Phe] (0.23)
JAK2 [p.Val617Phe] (0.66)
Available follow-up timepoints: 0-2-10-15-20
DNMT3A [p.Val830Ala] (0.72)
DNMT3A [p.Val830Ala] (1.72)
DNMT3A [p.Val830Ala] (3.04)
DNMT3A [p.Val830Ala] (2.96)
DNMT3A [p.Val830Ala] (2.44)
DNMT3A [p.Tyr660Cys] (0.08)
DNMT3A [p.Tyr660Cys] (0.17)
DNMT3A [p.Tyr660Cys] (0.4)
DNMT3A [p.Tyr660Cys] (0.96)
DNMT3A [p.Tyr660Cys] (1.29)
DNMT3A [p.Cys818Arg] (0.53)
TET2 [p.Arg571Ter] (0.12)
TET2 [p.Arg571Ter] (0.06)
Available follow-up timepoints: 0-10-15
DNMT3A [p.Gly646Arg] (2.33)
DNMT3A [p.Gly646Arg] (2.59)
DNMT3A [p.Gly646Arg] (0.97)
DNMT3A [p.Arg729Trp] (1.29)
DNMT3A [p.Arg729Trp] (1.28)
DNMT3A [p.Arg729Trp] (1)
NRAS [p.Ile21Val] (0.06)
NRAS [p.Ile21Val] (0.5)
NRAS [p.Ile21Val] (0.19)
GNAS [p.Cys843Arg] (0.3)
GNAS [p.Cys843Arg] (0.21)
TP53 [p.Arg174Gly] (0.12)
TP53 [p.Arg174Gly] (0.26)
TP53 [p.Arg174Gly] (0.34)

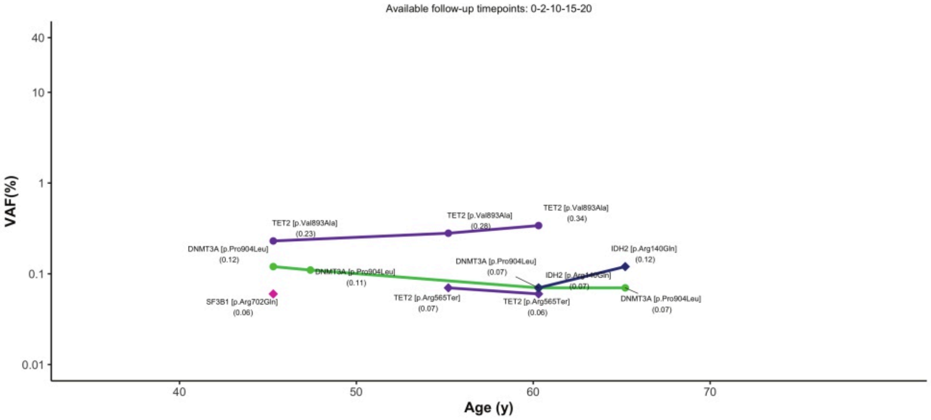
Available follow-up timepoints: 0-2-10-15-20
VAF(%)
Age (y)
TET2 [p.Val893Ala] (0.23)
TET2 [p.Val893Ala] (0.28)
TET2 [p.Val893Ala] (0.34)
DNMT3A [p.Pro904Leu] (0.12)
DNMT3A [p.Pro904Leu] (0.11)
DNMT3A [p.Pro904Leu] (0.07)
IDH2 [p.Arg140Gln] (0.12)
IDH2 [p.Arg140Gln] (0.07)
SF3B1 [p.Arg702Gln] (0.06)
TET2 [p.Arg565Ter] (0.07)
TET2 [p.Arg565Ter] (0.06)
DNMT3A [p.Pro904Leu] (0.07)

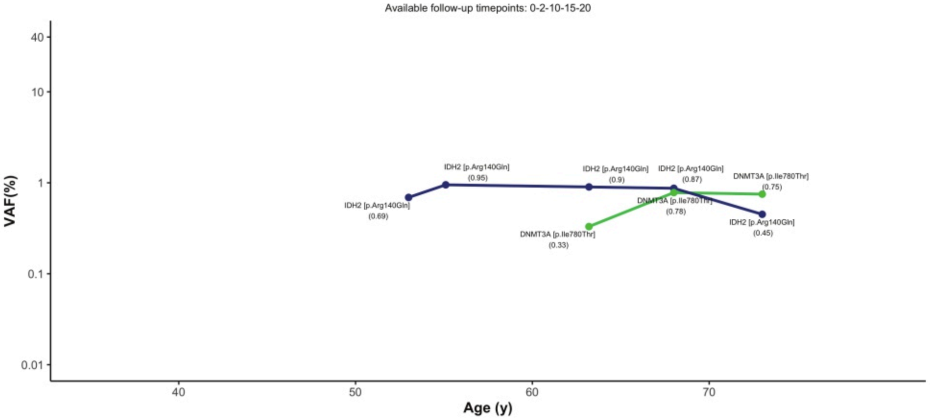
Available follow-up timepoints: 0-2-10-15-20
VAF(%)
Age (y)
IDH2 [p.Arg140Gln] (0.95)
IDH2 [p.Arg140Gln] (0.9)
IDH2 [p.Arg140Gln] (0.87)
DNMT3A [p.Ile780Thr] (0.75)
IDH2 [p.Arg140Gln] (0.69)
DNMT3A [p.Ile780Thr] (0.78)
IDH2 [p.Arg140Gln] (0.45)
DNMT3A [p.Ile780Thr] (0.33)

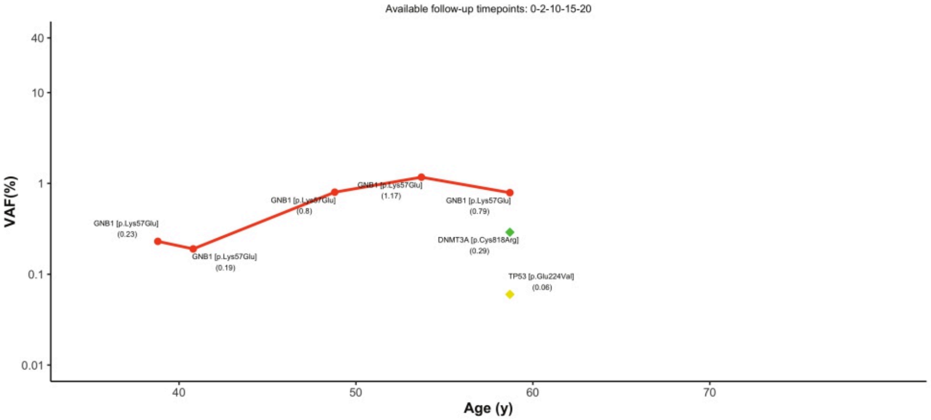
Available follow-up timepoints: 0-2-10-15-20
VAF(%)
Age (y)
GNB1 [p.Lys57Glu] (0.23)
GNB1 [p.Lys57Glu] (0.19)
GNB1 [p.Lys57Glu] (0.8)
GNB1 [p.Lys57Glu] (1.17)
GNB1 [p.Lys57Glu] (0.79)
DNMT3A [p.Cys818Arg] (0.29)
TP53 [p.Glu224Val] (0.06)

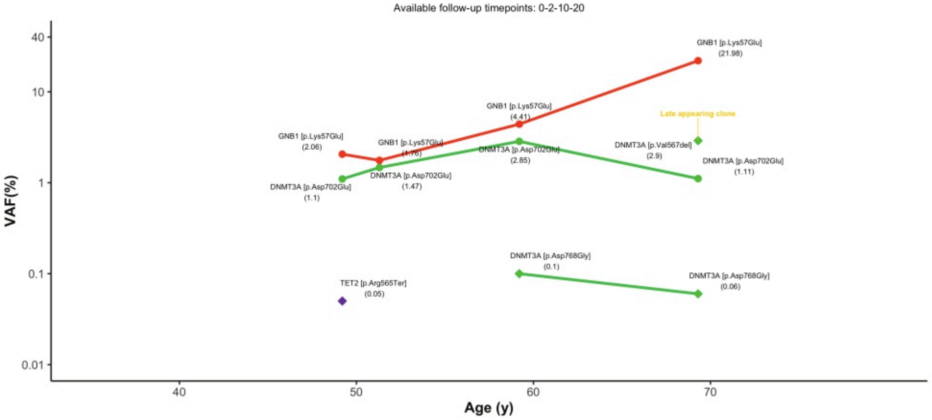
Available follow-up timepoints: 0-2-10-20
VAF(%)
Age (y)
GNB1 [p.Lys57Glu] (21.98)
GNB1 [p.Lys57Glu] (4.41)
Late appearing clone
GNB1 [p.Lys57Glu] (2.06)
GNB1 [p.Lys57Glu] (1.76)
DNMT3A [p.Asp702Glu] (2.85)
DNMT3A [p.Val567del] (2.9)
DNMT3A [p.Asp702Glu] (1.11)
DNMT3A [p.Asp702Glu] (1.1)
DNMT3A [p.Asp702Glu] (1.47)
DNMT3A [p.Asp768Gly] (0.1)
DNMT3A [p.Asp768Gly] (0.06)
TET2 [p.Arg565Ter] (0.05)

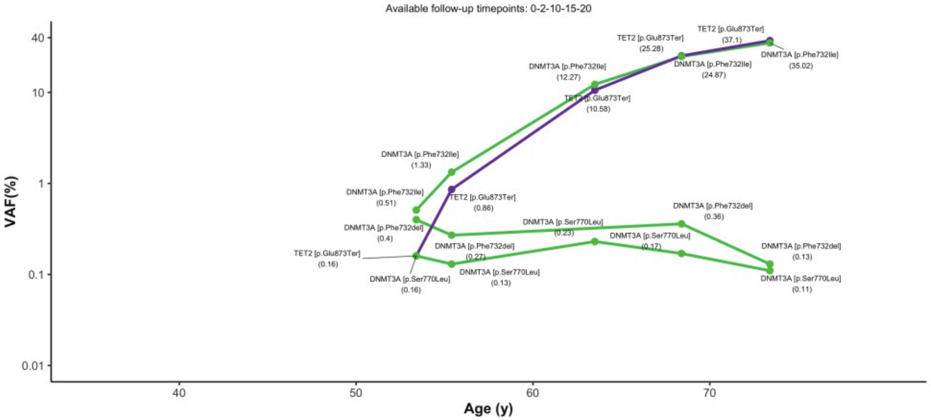
Available follow-up timepoints: 0-2-10-15-20
VAF(%)
Age (y)
TET2 [p.Glu873Ter] (37.1)
TET2 [p.Glu873Ter] (25.28)
DNMT3A [p.Phe732Ile] (35.02)
DNMT3A [p.Phe732Ile] (24.87)
DNMT3A [p.Phe732Ile] (12.27)
TET2 [p.Glu873Ter] (10.58)
DNMT3A [p.Phe732Ile] (1.33)
DNMT3A [p.Phe732Ile] (0.51)
TET2 [p.Glu873Ter] (0.86)
DNMT3A [p.Ser770Leu] (0.23)
DNMT3A [p.Phe732del] (0.36)
DNMT3A [p.Phe732del] (0.4)
DNMT3A [p.Ser770Leu] (0.17)
DNMT3A [p.Phe732del] (0.27)
DNMT3A [p.Phe732del] (0.13)
TET2 [p.Glu873Ter] (0.16)
DNMT3A [p.Ser770Leu] (0.16)
DNMT3A [p.Ser770Leu] (0.13)
DNMT3A [p.Ser770Leu] (0.11)

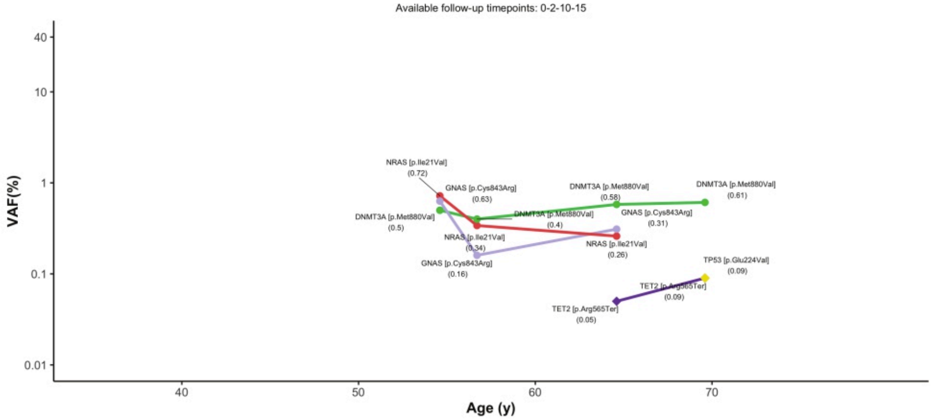
Available follow-up timepoints: 0-2-10-15
VAF(%)
Age (y)
NRAS [p.Ile21Val] (0.72)
GNAS [p.Cys843Arg] (0.63)
DNMT3A [p.Met880Val] (0.58)
DNMT3A [p.Met880Val] (0.61)
DNMT3A [p.Met880Val] (0.5)
DNMT3A [p.Met880Val] (0.4)
GNAS [p.Cys843Arg] (0.31)
NRAS [p.Ile21Val] (0.34)
NRAS [p.Ile21Val] (0.26)
GNAS [p.Cys843Arg] (0.16)
TP53 [p.Glu224Val] (0.09)
TET2 [p.Arg565Ter] (0.09)
TET2 [p.Arg565Ter] (0.05)

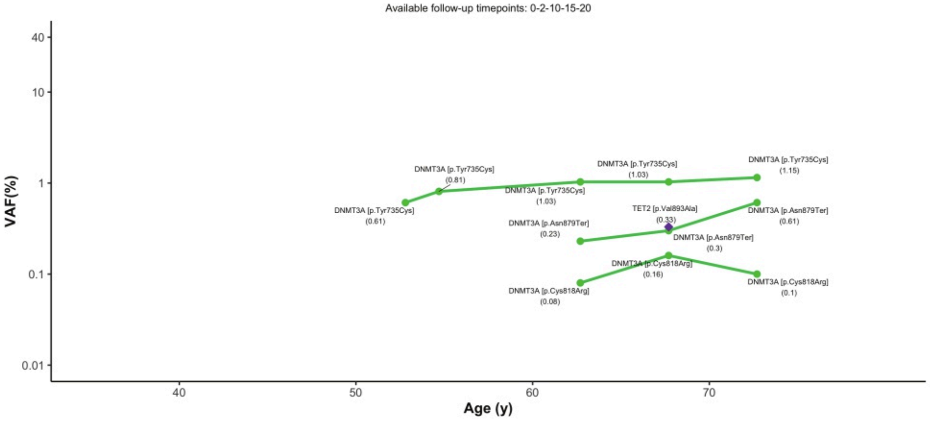
Available follow-up timepoints: 0-2-10-15-20
VAF(%)
Age (y)
DNMT3A [p.Tyr735Cys] (0.81)
DNMT3A [p.Tyr735Cys] (1.03)
DNMT3A [p.Tyr735Cys] (1.15)
DNMT3A [p.Tyr735Cys] (1.03)
DNMT3A [p.Tyr735Cys] (0.61)
TET2 [p.Val893Ala] (0.33)
DNMT3A [p.Asn879Ter] (0.23)
DNMT3A [p.Asn879Ter] (0.61)
DNMT3A [p.Asn879Ter] (0.3)
DNMT3A [p.Cys818Arg] (0.16)
DNMT3A [p.Cys818Arg] (0.08)
DNMT3A [p.Cys818Arg] (0.1)

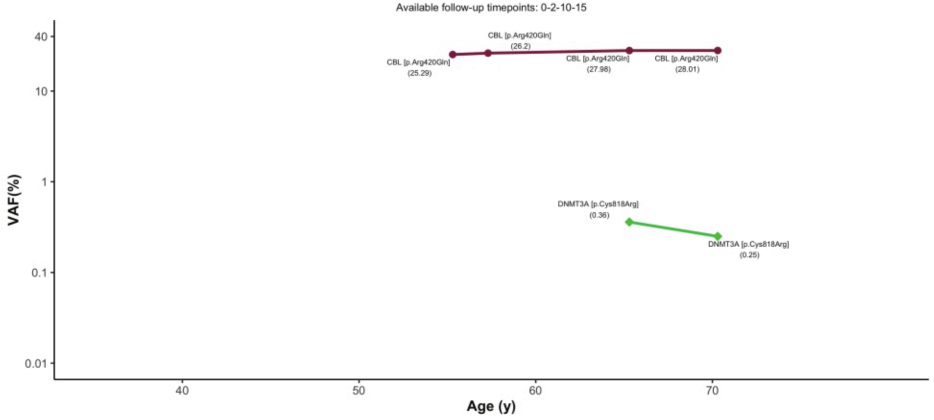
Available follow-up timepoints: 0-2-10-15
VAF(%)
Age (y)
CBL [p.Arg420Gln] (26.2)
CBL [p.Arg420Gln] (25.29)
CBL [p.Arg420Gln] (27.98)
CBL [p.Arg420Gln] (28.01)
DNMT3A [p.Cys818Arg] (0.36)
DNMT3A [p.Cys818Arg] (0.25)

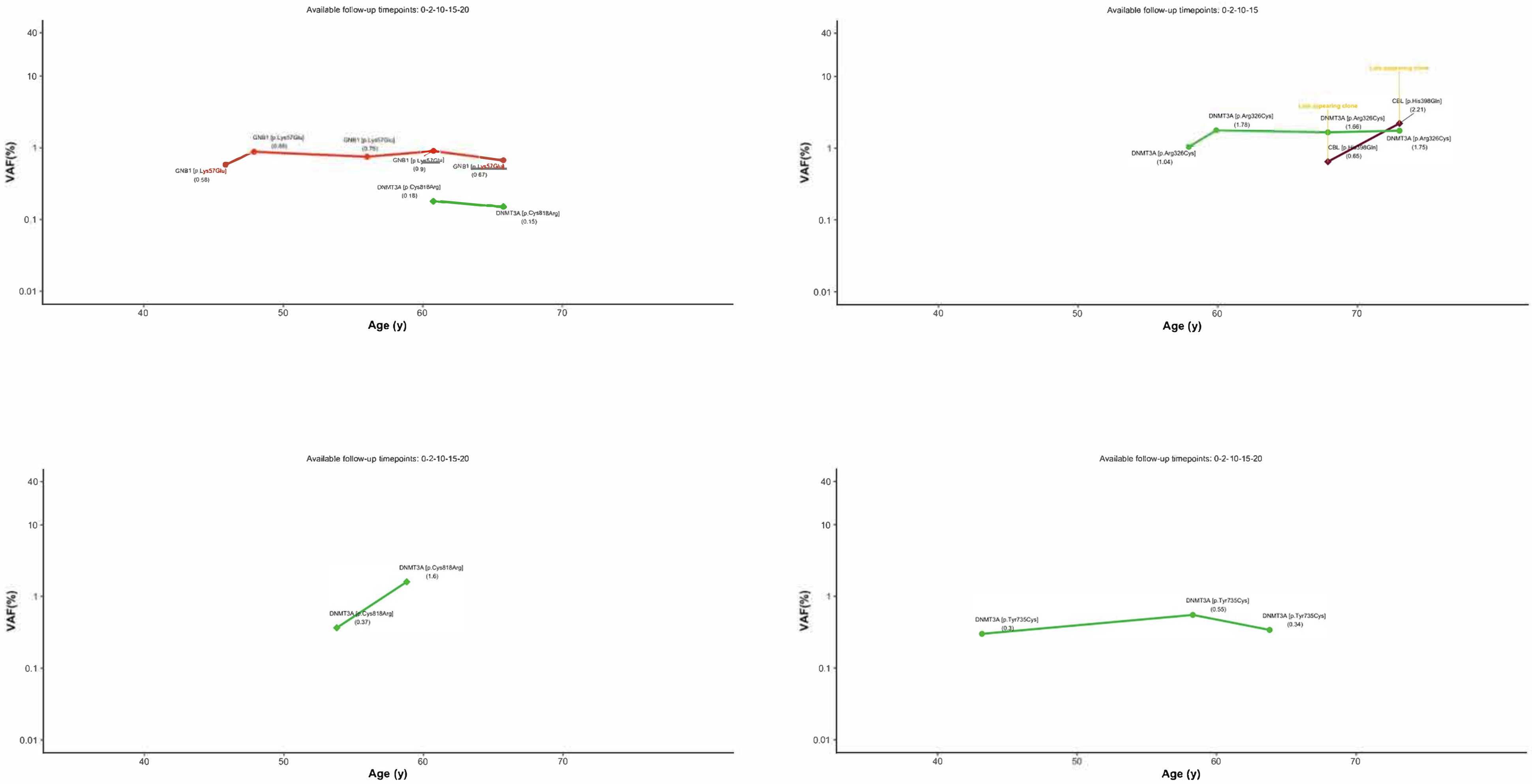


***File S6.*** **Individual plots of CHDMs detected during follow-up time**. Each graph represents one individual. Each point is labelled with the gene name, Amino Acid reference residue, -position, and -alternative residue, with the VAF of the CHDM at that point below in brackets. Late appearing clones are annotated with an additional label "late appearing clone". CHDMs categorized as events are shown in triangles, trajectories in circles.
