## Supplementary FileS7 (figure 5 in supplement) for "Expansion of mutation-driven haematopoietic clones is associated with insulin resistance and low HDL-cholesterol in individuals with obesity"

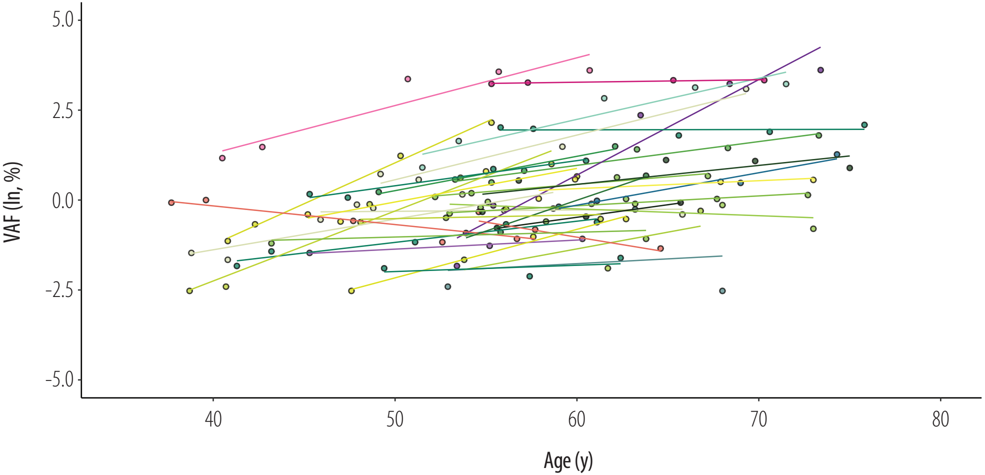


***File S7*. Slope inspection of 32 selected trajectories in multiple-timepoint dataset.** A linear regression line as computed by geom_smooth() is shown for each selected trajectory.
