## Supplementary FileS8 (figure 6 in supplement) for "Expansion of mutation-driven haematopoietic clones is associated with insulin resistance and low HDL-cholesterol in individuals with obesity"

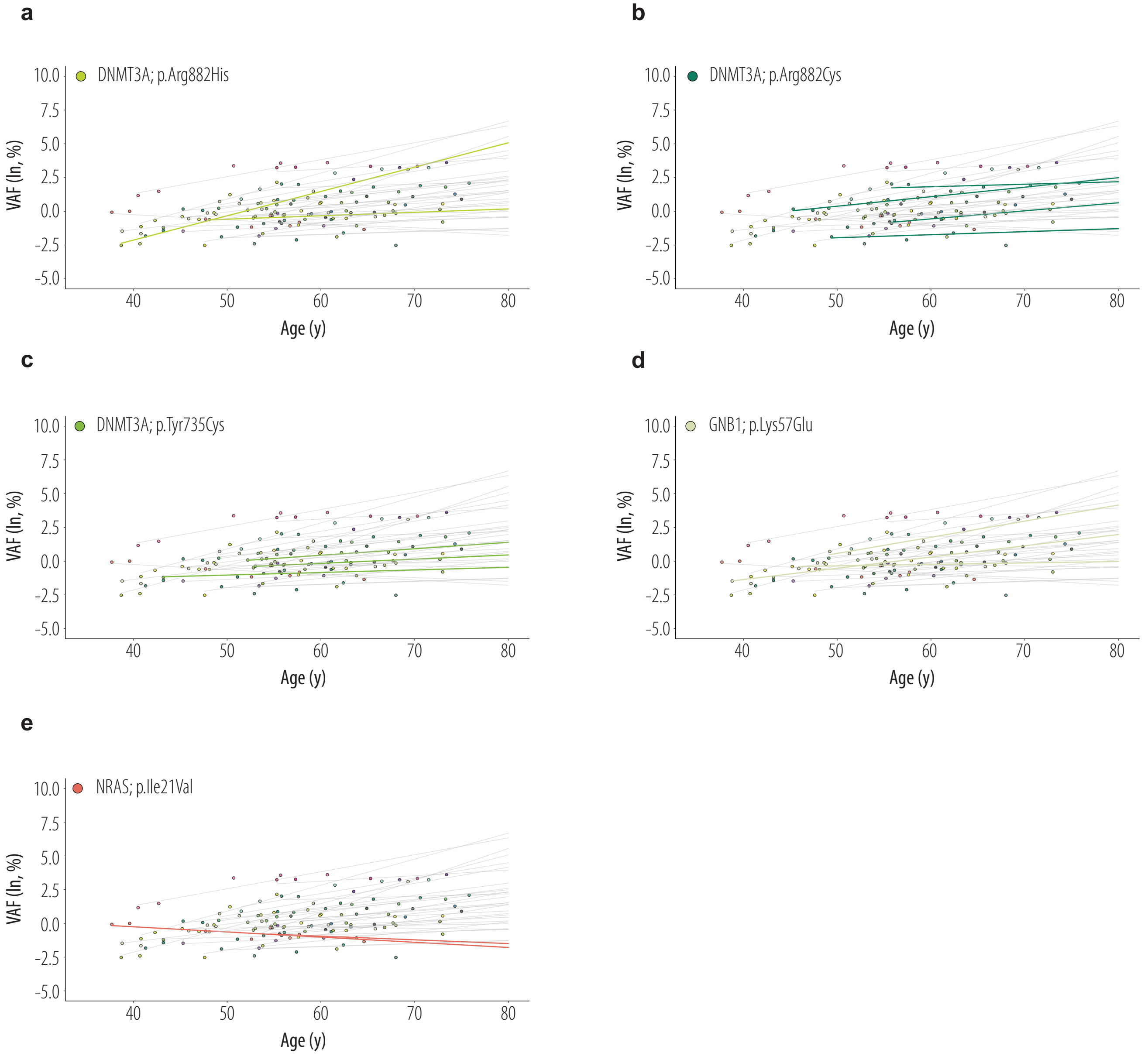


*File S8.* **Difference in rate of growth for recurrent trajectories in independent individuals. (a)** The trajectory DNMT3A;p.Arg882His was identified in two different individuals as the most important trajectory. **(b)** The trajectory DNMT3A;p.Arg882Cys was identified in four different individuals as the most important trajectory. **(c)** The trajectory DNMT3A;p.Tyr735Cys was identified in three different individuals as the most important trajectory. **(d)** The trajectory GNB1;p.Lys57Glu was identified in three different individuals as the most important trajectory. **(e)** The trajectory NRAS;p.Ile21Val was identified in two different individuals as the most important trajectory.
